# The WRC domain of GRF transcription factors. Structure and DNA recognition

**DOI:** 10.1101/2025.01.03.631214

**Authors:** Franco A. Biglione, Javier F. Palatnik, Nahuel D. González Schain, Rodolfo M. Rasia

**Affiliations:** Instituto de Biología Molecular y Celular de Rosario (IBR-CONICET-UNR), Rosario 2000, Santa Fe, Argentina; Área Biofísica, Facultad de Ciencias Bioquímicas y Farmacéuticas, Universidad Nacional de Rosario, Rosario 2000, Santa Fe, Argentina

## Abstract

Growth-regulating factors (GRFs) belong to a plant-specific family of transcription factors which display important roles in plant growth and development. GRF transcriptional activity is finely tuned by regulatory processes involving post-transcriptional silencing exerted by miRNA396, and protein-protein interactions involving a family of co-transcriptional regulators known as GRF-interacting factors (GIFs). In this way, expression of GRF target genes is modulated by a highly complex interplay between GRF/GIF isoform diversity and expression patterns along with miR396 and GIF gradients throughout plant tissues. At the protein level, GRFs are composed of two highly conserved domains known as QLQ and WRC and a less conserved C-terminal trans-activation domain. Whereas QLQ mediates GRF-GIF interaction by forming a complex with SNH domains found in GIFs, the WRC has been proposed as a putative zinc finger domain responsible for target DNA recognition and nuclear import. However, the structural aspects governing GRF transcriptional activity and target recognition remain unknown. In this work, we applied bioinformatic and biophysical analysis to comprehensively characterize the structural features that modulate the biological function of this protein family with a focus on the WRC domain. We provide insights into the structure of the WRC domain in GRFs and explore the WRC features driving GRFs:DNA complex formation. These findings offer new insights into how WRC domains modulate the biological functions of GRFs, laying the groundwork for future studies on their structure-function relationship in gene regulation and development of plants.

## 1 Introduction

Eukaryotic organisms rely on complex regulatory networks to proficiently perform their biological functions. The differential activity of genes is finely tuned by the action of transcription factors that orchestrate the expression of specific genes through binding to DNA cis-targeting sequences. Among these factors, GROWTH-REGULATING FACTORs (GRFs) play a crucial role in plant growth and development such as the regulation of cell proliferation in leaves ^1–4^. GRFs are widely conserved in land plants, in the form of gene families of between 8 and 20 members. For instance, *Arabidopsis thaliana*, rice (*Oryza sativa*), and soybean (*Glycine max*) possess 9, 12, and 24 GRF members, respectively ^5^. GRFs were shown to bind to regulatory sequences near target genes and modify their expression levels ^6–9^.

At the structural level, GRFs span between 250 and 500 residues in length and are defined by the presence of two conserved domains at their N-terminus: the QLQ and the WRC domains ^2^. The QLQ domain, which has 36 residues, contains a conserved QX_3_LX_2_Q motif and is involved in the interactions with the transcriptional coactivators of the GRF-INTERACTING FACTORS family (GIFs) ^3^. On the other hand, the presence of a strictly conserved CX_9_CX_10_CX_2_H motif indicates that the WRC domain is a putative zinc finger DNA-binding motif ^1,10^. Finally, the C-terminal region is highly variable in length and composition and is associated with the transactivation activity of GRF proteins ^3,11^.

The regulatory role of GRFs in proliferating cells extends beyond leaves to encompass other organs in a wide variety of plant species. In this sense, these transcription factors, together with the GIF coactivators, participate not only in the growth of leaves but also roots, stems and inflorescences. Moreover, they also influence tolerance to UV light and drought, and lead to delayed senescence in Arabidopsis. GRF expression and localization is in turn subject to post-transcriptional regulation by miR396, whose recognition site is found within the coding sequence of the WRC domain ^12^.

Central to their function, GRFs interact with specific *cis* regulatory sequences of different genes ^6–8,13^. Deletion of the WRC domain of barley GRF1 (BGRF1) prevents binding of the protein to the promoter region of the Bkn3 gene ^6^. It has also been shown that *A. thaliana* GRF7 (AtGRF7) controls the transcription of the transcriptional activator DREB2A by binding to a *cis* regulatory element (TGTCAGG) in its promoter ^7^. In particular, GRF7 and GRF10 from *Oryza sativa* (OsGRF7/10) modulate the expression of the rice KNOX Oskn2 gene, involved in the regulation of meristematic function, through the interaction with a 34 bp fragment in the promoter region ^8^. Interestingly, although this region lacks the TGTCAGG sequence, it is also recognized by AtGRF4, AtGRF5, and AtGRF6, and binding requires both the WRC and QLQ domains. This finding suggests that the specificity of WRCs for DNA sequences may be relaxed. In addition, a ChIP-seq study of AtGRF1 and AtGRF3 resulted in enrichment in the ACTCGAC and CTTCTTC sequences ^13^. These divergent results challenge our understanding of the specificity of WRC domains for DNA sequences. Moreover, the residues responsible for DNA recognition within this domain remain unknown.

The WRC domain has been classified as a non canonical Zn(II) finger domain due to its similarity to the HRT (Hordeum Repressor of Transcription) protein found in barley ^10^, yet it has not been experimentally tested. Unlike canonical Zn fingers, the conserved pattern in the WRC domains is C-X_7_-WRC-X_10_-C-X_2_-H. Its classification as a Zn finger responds to the conservation of the 4 putative metal ligand residues (CCCH), but its structural properties remain unknown due to the lack of homology to any Zn finger domain of known structure. Zinc finger domains display extensive sequence and structural variability and the metal coordination is essential for their folding ^14^. While critical for structural integrity, the Zn(II) ion typically does not directly engage in DNA binding. Although most of the characterized Zn fingers adopt a ββα fold, there are numerous variants that accommodate different numbers of residues between the metal ligands giving rise to remarkably diverse structural arrangements ^15^. The coordination of the metal ion offers a strong stabilization compared to other short domains, and gives it the possibility to explore a wide sequence space, enhancing its versatility.

The WRC domain sequence, spanning only 44 residues, is strictly found in plants and experiences significant evolutionary pressure within GRFs and potentially other proteins containing this domain. It features a putative bipartite nuclear localization signal composed of basic residues (R/K) at 11 specific positions along its sequence. Additionally, the domain coding sequence harbors a miR396 binding site critical for post-transcriptional regulation and proper plant development ^16^. Despite its role in binding to DNA and guiding the GRFs to selectively regulate gene expression, the residues within the WRC domain responsible for the recognition of GRF cis-targeting sequences remain elusive. Delineating the mechanisms by which this domain interacts with DNA is essential to understand differences in target gene specificity for WRC-containing proteins, both within and outside the GRF family of transcription factors.

The WRC domain, originally identified in Growth-Regulating Factors (GRFs), is not confined solely to this family of transcription factors but has also been discovered in other plant proteins. A recent study described that JMJ28, a Jumonji C domain-containing protein in *Arabidopsis*, harbors a WRC domain which is essential for directing ATX1/2-containing COMPASS-like complexes to specific chromatin targets ^17^. However, this domain’s presence across diverse protein architectures ^18^ and its sequence heterogeneity has not yet been explored.

In this study, we conducted bioinformatic and structural analysis of WRC domains specifically focusing on those from *Arabidopsis thaliana* GRF1, GRF3, GRF5, and GRF7. We found that all these domains bind one Zn(II) ion per monomer with high affinity, which is crucial for their proper folding. The folded proteins exhibit moderate stability, with melting temperatures ranging from 38°C to 56°C. Finally, our characterization revealed that these GRFs can bind dsDNA in a nonspecific fashion but display preferences for their specific cognate sequences.

## 2 Methods

### 2.1 Bioinformatic analysis

WRC-containing proteins were downloaded from UniProt using the PFAM ID (PF08879). Briefly, the WRC sequences were curated by extracting WRC extended sequences from each full-length primary sequence, extending beyond the WRC limits reported by UniProt by 25 amino acids at both ends, when possible. These extended sequences were then aligned using MAFFT. Conservation analysis within the alignment facilitated the proper determination of WRC boundaries, which were further trimmed by removing non-conserved terminal regions to obtain more compact WRC sequences. Sequences with >90% identity were discarded, along with those shorter or longer than the median sequence length by more than 30%. The length-filtered set was then re-aligned, and positions containing gap residues in more than 50% of the sequences were eliminated (Figure S1A). SVD analysis was performed using previously established protocols and open-source Python scripts.

### 2.2 Plasmid construction

The *A. thaliana* WRC sequences from GRF1, GRF3, GRF5, and GRF7 (UniProt accession numbers O81001, Q9SJR5, Q8L8A6, and Q9FJB8, respectively) were amplified from cDNA samples (Table S3) by PCR and inserted into the pET-32a-derived pT7 expression vector using restriction-free cloning ^19^. This vector provides a His tag, followed by the TrxA fusion protein and a TEV cleavage site at the 5’ end of the target gene. Constructs were verified by DNA sequencing. All expressed WRC domains span 5 amino acids beyond the reported limits of the WRC sequences for these GRF isoforms at both ends. All DNA oligonucleotides and sequencing services were obtained from Macrogen (Seoul, Republic of Korea).

### 2.3 Protein expression and purification

Expression plasmids were transformed in *E. coli BL21(DE3)* cells which were then grown at 37 °C in Erlenmeyer flasks shaken at 180 rpm in either M9 minimal medium supplemented with 1 g/L ^15^N-NH_4_Cl or 1 g/L ^15^N-NH_4_Cl and 2 g/L U[^13^C]-Glucose (Cambridge Isotope Laboratories), or in LB broth in the presence of 100 μg/mL Ampicillin. Protein expression was induced with 0.25 mM IPTG at OD600 ≈ 0.7 and cells were incubated for an additional 4 h at 37°C. The cells were collected by centrifugation at 4000 g for 20 min at 4 °C, followed by resuspension into a 20 mL solution containing 50 mM Tris pH 8.0, 500 mM NaCl, 1 mM β-mercaptoethanol. The suspension was disrupted by sonication and the soluble fraction was clarified by centrifugation for 30 min at 20,000 g. His-tagged Trx-WRC fusions were purified using a Ni(II) column and digested with His-tagged TEV protease. The digested proteins were purified with a Superdex 75 Increase 10/300 GL size exclusion chromatography column equilibrated with 20 mM HEPES, 500 mM NaCl, pH 7.0. Proteins were concentrated using centrifugal filter units and supplemented with TCEP 5 mM and 1 equivalent of ZnSO_4_. Protein concentration was measured by UV absorption at 280 nm using the corresponding absorptivity coefficient calculated by ProtParam tool at ExPASy web portal ^20^. Sequences of purified WRC variants are detailed in Table S4. WRC samples were used directly or flash-frozen in liquid nitrogen and transferred immediately to a -80°C freezer for long-term storage.

### 2.4 Zn(II) stoichiometry and binding affinity

The zinc(II) content was determined using the metallochromic indicator 4-(2-pyridylazo)resorcinol (PAR) ^21^. HoloWRC7 protein samples (6.2 µM) were incubated with PAR (100 µM) in a denaturing buffer (2.8 M guanidinium chloride, 20 mM HEPES, and 50 mM NaCl, pH 7.0), at 100°C for 20 minutes to release bound zinc. Absorbance was measured at 500 nm, with background correction at 650 nm, using a 96-well plate. Zinc concentrations were quantified against a Zn²⁺ calibration curve (0–10 µM) prepared from a ZnCl₂ standard solution. The results represent the average of triplicate measurements. All buffer solutions were treated with Chelex 100 (Sigma) by extensive stirring to remove trace metal contamination. Prior to zinc quantification, HoloWRC7 was diafiltered into Chelex-treated buffer (20 mM HEPES, 50 mM NaCl, pH 7.0) to remove unbound zinc and minimize interference. A protein-free buffer was used as a control for quantification.

ApoWRC7 was prepared by acidifying WRC7 in 20 mM HEPES, 50 mM NaCl, pH 2.0, at 4°C for 4 hours to remove bound zinc. The sample was subsequently concentrated and subjected to three cycles of diafiltration in Chelex-treated buffer at pH 2.0 to remove residual metals. During this process, 5 mM TCEP was added to prevent cysteine oxidation. The buffer was then exchanged to 20 mM HEPES, 50 mM NaCl, and 5 mM TCEP at pH 7.0. Dissociation constants for Zn(II) binding were estimated by competition with the chromophoric chelator PAR ^21^. PAR is a metallochromic compound whose UV-visible absorption spectrum changes upon metal coordination, resulting in a shift of its maximum absorption wavelength from 414 nm to 500 nm. Pure PAR and PAR_2_Zn spectra were used to deconvolute the spectra of each species using Multivariate Curve Resolution-Alternating Least Squares (MCR-ALS). The dissociation constant of PAR_2_-Zn under our experimental conditions was estimated as 2.2 10^-12^ M^2^, consistent with previously reported values at pH 7 ^21^. This value was then used to calculate the Kd of WRC-Zn using the same deconvolution procedure. The binding curve of PAR competition experiments was fitted to a one-site binding model for WRC7, as measured by our Zn^2+^ content determination experiments. Spectra were measured at 25°C using a Jasco V-630 BIO spectrophotometer (Jasco, Easton, MD, USA) with Peltier temperature control (10 mm quartz cell) and each spectra corresponds to an independently prepared sample. All Kd fittings were performed by using the DynaFit software package ^22^.

### 2.5 Circular dichroism spectroscopy

Circular dichroism (CD) spectra were recorded using a Jasco J-1500 spectropolarimeter (Jasco, Easton, MD, USA).

For far-UV CD measurements, protein samples were placed in a 1 mm pathlength quartz cuvette to minimize buffer absorption. Spectra were recorded from 190 to 240 nm at 10°C, averaging four scans to enhance the signal-to-noise ratio. The buffer contained 50 mM phosphate at pH 7.0, and the final protein concentration was 10–20 μM (∼0.1 mg/mL). Samples were freshly prepared by diluting stock WRC solutions (Section 3.2).

Near-UV CD spectra were recorded from 250 to 320 nm at 10°C using a 10 mm pathlength quartz cuvette. Protein samples were prepared at a final concentration of 120–200 μM (A₂₈₀ ≈ 1) in the stock buffer. Four scans were averaged for each spectrum.

To obtain CD spectra for ApoWRC, 10 molar equivalents of EDTA were added to chelate zinc, ensuring complete demetallation.

Thermal melting experiments were performed by recording near-UV spectra from 10°C to 95°C in 5°C increments, with a heating rate of 1°C/min. The data sets of melting spectra were analyzed using MCR-ALS to resolve the spectra of folded and unfolded states. Melting temperatures (Till) were determined by fitting the evolution of the folded component as a function of temperature to a two-state transition model.

CD data were processed using an in-house developed library specifically designed for reading and processing CD spectra (github.com/francobiglione/ProteinBiophysics), along with custom Python scripts for data analysis and visualization.

### 2.6 NMR spectroscopy

HSQC NMR spectra of ^15^N labelled WRCs were acquired at 298 K on a 700 MHz Bruker Avance III spectrometer equipped with a TXI probehead using pulse sequences from the standard Bruker library. All spectra were processed with NMRPipe and analyzed with CCPNMR in the NMRBox software suite ^23–25^. ApoWRC forms were generated as described in section 3.5.

The WRC3 construct was assigned using a set of triple-resonance spectra (HNCA/(CO)CA, HNCACB/(CO)CACB, HN(CA)CO/HNCO) and ^1^H-^15^N heteronuclear TOCSY and NOESY datasets, collected on the same spectrometer for double (^13^C-^15^N) or simple ( ^15^N) uniformly labeled samples at 298 K and stock buffer conditions with 10% D_2_O (section 3.3). Secondary structure populations were predicted by implementation of the simultaneous sequence-based predictor s2D by means of assigned backbone chemical shifts (H_α_,CO,C_α_,C_β_,H_N_,N_H_) ^26^. Amide hydrogen bond interactions were assessed by calculating temperature coefficients derived from ^15^N-^1^H HSQC spectra recorded at 298, 303, 308, 313 and 318 K ^27^. Similarly, 2,2,2-trifluoroethanol (TFE) coefficients were calculated from ^5^N-^1^H HSQC recorded at 298 K at increasing ratios of TFE/buffer mixtures (0, 4, 9 and 15 % v/v). These coefficients were used to map secondary structure elements based on TFE’s ability to perturb them ^28^. For temperature and TFE coefficients, error bars in the plots represent standard errors for the linear fit of each residue position. HN-N Residual dipolar couplings (RDCs) were measured at 298 K in C12E5 n-hexanol anisotropic medium after stabilization of the sterically induced alignment ^29^. HN-N RDCs were obtained on a ^15^N labelled sample using the standard IPAP sequence, as implemented in the Bruker library. Analysis and calculations of RDCs were performed with the best-ranked AlphaFold (AF3) WRC3 model using the software MODULE2 ^30^. Backbone dynamics of WRC3 were determined at 298 K. ^15^N T1 and T2 relaxation and ^1^H-^15^N nuclear Overhauser effect (NOE) experiments were recorded using standard pulse sequences from the Bruker Topspin library. For ^15^N T1 and T2 acquisitions, relaxation delays were randomized, and duplicate spectra were collected at several time points to estimate uncertainties. Relaxation rates were calculated by fitting peak intensities at different time points to a two-parameter exponential decay function using CCPNMR. Steady-state ^1^H-^15^N hetNOE values were obtained by dividing the peak heights of paired spectra collected with and without an initial 4-second proton saturation period. Correlation time (τ_c_) was calculated from the R2/R1 ratio as described elsewhere ^31^. Tryptophan and histidine sidechain resonances were assigned using homonuclear ^1^H-^1^H TOCSY and NOESY (D_2_O) experiments and long-range ^15^N-^1^H HSQC spectra acquired at 278 K on non-labelled WRC7 or ^15^N WRC7, respectively, using pulse sequences from the standard Bruker library. NOESY samples were generated by diafiltration with D_2_O-prepared stock buffer. All illustrations containing NMR spectra were generated using Python’s standard visualization libraries and the nmrglue module ^32^.

### 2.7 DNA interaction

#### Probes for DNA Interaction Studies

Probes used in DNA interaction studies are detailed in Table S3. For fluorescence and NMR experiments, we utilized specific and non-specific GRF7 long oligonucleotides (34 bp), previously labeled as E1 and E2 in the original publication ^7^. For CD and NMR studies, minimal specific dsDNA sequences were designed based on reported GRF1 (13 bp) and GRF7 (12 bp) cis-targeting sites. A minimal non-specific probe, derived from a nucleotide region shown to lack GRF7 interaction ^7^, was used for GRF1 NMR studies due to the similarity of cis-targeting sites between GRF7 (T<u>GTCAGG</u>) and GRF1 (reverse complement of <u>GTC</u>G<u>AG</u>T*). Minimal probes were designed as tetraloops to ensure the dsDNA structure at GRF-targeting sites was maintained, minimizing the presence of interfering ssDNA. Loop nucleotide identity and closing base pairs were rationally selected to optimize tetraloop stability based on well-characterized studies in the literature ^33^. Long probes were annealed to dsDNA by heating stock solutions at 95 °C for 5 minutes, followed by slow cooling to 20 °C. Minimal dsDNA was prepared by heating stock solutions for 5 minutes at 95 °C, then snap cooling in an ice bath to prevent intermolecular interactions.

#### Fluorescence Binding Studies

Fluorescence equilibrium binding isotherms for WRC7 were obtained by monitoring tryptophan quenching upon nucleic acid binding. Fluorescence measurements were conducted using a Cary Eclipse spectrofluorometer with an excitation wavelength of 280 nm and emission spectra recorded from 290 to 450 nm. Data were collected at 15 °C, following the addition of 15 µM long or minimal GRF7 oligonucleotide solutions to 50 µL protein samples (1 µM) at varying DNA/protein ratios (0–2 equivalents). The buffer conditions were 20 mM HEPES, 50 mM NaCl, pH 7.0. Each fluorescence spectrum corresponded to an independent sample of a defined DNA/WRC ratio, measured in a 50 µL quartz cell. Spectra were corrected for baseline contributions from buffer fluorescence. WRC7-nucleic acid complex formation was quantified by the reduction in initial protein fluorescence, expressed as fraction quenching (1−*F*/*F*_0_), and plotted against bp/WRC equivalents (DNA/WRC ratio multiplied by oligonucleotide length).

#### Circular Dichroism (CD) Binding Studies

CD spectra for DNA-binding analyses were recorded from 190 to 320 nm at 10 °C , averaging four scans to enhance signal-to-noise ratio. The buffer condition was 50 mM phosphate, pH 7.0. Protein concentrations were 13 µM, and minimal dsDNA (30 µM stock solutions) were added at DNA/WRC ratios ranging from 0 to 2 equivalents. Difference CD spectra were generated by subtracting blank DNA spectra at the same oligonucleotide concentrations. Each protein-DNA or blank DNA spectrum was measured from an independent sample. Binding signals were derived by integrating the area under the curve (270–300 nm) using Riemann’s approximation. bp/WRC equivalents were calculated as described previously. To assess the lack of binding of ApoWRC1 to WRC1 minimal sequences, circular dichroism (CD) measurements were performed in the presence of 0.5 equivalents of minimal dsDNA probes and 1.6 mM EDTA to chelate zinc ions.

#### NMR chemical shift perturbation analysis

^15^N-^1^H HSQC NMR spectra of WRC1:dsDNA samples were acquired at 298 K (section 3.6) in 3 mm NMR tubes. Mock buffer, minimal specific or minimal nonspecific DNA probes (0.9-1 mM stock DNA solutions) were stepwise added at 0.5 and 1 equivalents to three independent ^15^N WRC1 samples before acquisition of spectra at these conditions (samples were diluted in stock buffer). WRC1 assignment was performed by transferring WRC3 assignment of conserved residues onto WRC1 peaks of spectra without DNA, signals for which there was no clear assignment were removed. Chemical shift displacement was identified by analysing disappearing signals and appearing of new signals nearby and confirmed through the analysis of spectra at the two increasing concentrations to account for signal direction of displacements. Quantification and mapping of DNA-binding residues was performed by calculation of chemical shift perturbations of each residue using the following equation: 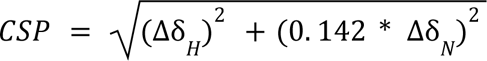. Mean CSP plus half standard deviation was used as a threshold for identifying perturbed residues involved in DNA binding.

## 3 Results

### 3.1 WRC domains from GRFs share a distinct amino acidic imprinting

To gain insight into the uniqueness of WRC domains present in GRFs among the WRC-containing proteins, we performed singular value decomposition (SVD) of WRC sequences belonging to heterogeneous protein architectures. With this aim, 6957 protein sequences with an associated WRC PFAM (PF08879) were retrieved from UNIPROT, all of which represented 2083 non-duplicated WRC sequences. The unique WRCs were further filtered based on their sequence identity (<90%) and length (60% interval around the median length) to reduce redundancy and retain sequence variability (Figure S1A). A total of 1244 curated WRCs were used to generate a multiple sequence alignment (MSA) which was finally analyzed through SVD ^34^. The nearly linear correlation of the consensus identity of WRC sequences in the MSA and σ_1_u_i_^(1)^ with a pearson coefficient of 0.989 indicates a strong relationship between conservation and the first singular coordinate, implying that the dataset proficiently captures overall residue conservation (Figure S1B). The consensus sequence for the entire dataset suggests a resemblance to WRC sequences found in GRFs (Figure S1D).

The projection of sequences in SVD space revealed that WRCs are not randomly distributed but instead cluster into distinct groups, suggesting they may belong to protein subfamilies with divergent functions (Figure 1B). Clustering using K-means resulted in an optimal of 3 clusters which are well resolved in the first three singular coordinates (Figure S1C). Surprisingly, we found that WRC-containing proteins from *A. thaliana* segregate based on their biological function. A detailed cluster-specific analysis of the proteins harboring the WRC sequences showed that each SVD subgroup displays preferential enrichment for specific domain architectures in which the WRC domain is embedded (Table S1), being GRFs almost exclusively grouped into a distinct cluster (Table S2).

**Figure 1.**
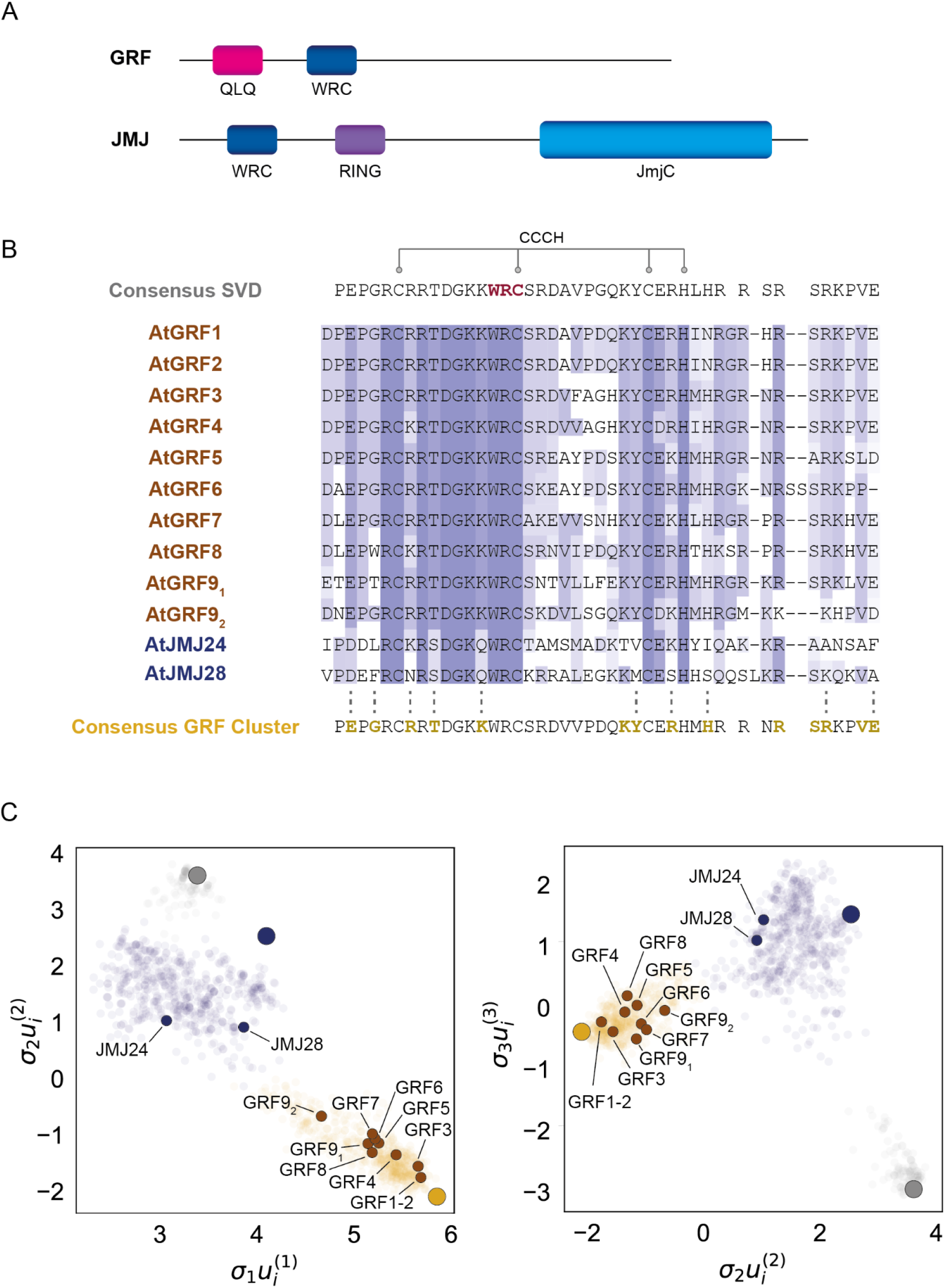
Singular Value Decomposition analysis of WRC sequences. (A) Domain architecture of WRC-containing proteins. (B) Alignment of A. thaliana WRC-containing proteins and comparison to consensus sequences derived from the analysis for the whole sequence dataset (Consensus SVD, top) or the cluster containing WRC domains from GRFs (Consensus GRF cluster, bottom). The alignment is colored according to the BLOSUM62 score with a conservation threshold of 16%. Highlighted residues on the consensus SVD sequence correspond to the WRC motif (bold) and zinc binding residues (CCCH motif). Residues with a preferential in-cluster vs out-of-cluster enrichment of at least 60% for the consensus GRF cluster are highlighted in yellow. Dashed lines indicate enriched consensus GRF cluster residues not present in JMJ proteins. (C) Mapping of A. thaliana WRC-containing proteins along the clusters identified in the first three singular coordinates. Larger circles denote the coordinates of the consensus sequences calculated for each cluster.

Segregation of WRC from GRFs into a specific cluster in the SVD space suggests the presence of intrinsic features central to their function that are encoded in the domain sequence. To characterize the relevant residues responsible for cluster partitioning we calculated the consensus sequences of each cluster and compared the frequency of each residue at each position between different clusters (Figure S1D). In the case of the cluster that includes WRCs from proteins with a GRF architecture, the residues having a preferential enrichment are in good agreement with the sequence differences found between AtGRFs and AtJMJ24/AtJMJ28 at the level of the WRC domain (Figure 1B). As a result, our SVD analysis allowed us to identify potential WRC residues that may define the GRF nature of the WRC domains within the specific architecture of this family of transcription factors.

### 3.2 The WRC is a zinc finger domain that folds in the presence of Zn^2+^

We next carried out structural studies to properly assess the role of the conserved residues within the domain in atomic detail. As shown in Figure 1A, *Arabidopsis thaliana* has nine GRF genes which have been reported to belong to five different phylogenetic groups ^5^. We selected one representative sequence from four phylogenetic groups, leaving out the group of AtGRF9 as this isoform has been suggested to function differently from other GRFs due to the presence of two WRC domains ^35^. Therefore we expressed the WRC domains from AtGRF1, AtGRF3, AtGRF5, AtGRF7 as C-terminal fusions to TrxA. All constructs were soluble, and the corresponding WRC domains were obtained by digestion of the fusion products.

We studied the overall folding properties of the four isoforms using circular dichroism. The near-UV spectra of all domains show that they have a stable tertiary structure with well-structured bands (Figure 2A and Figure S2A). On its side the far-UV spectra are less well defined, indicating that the domains have little content of canonical secondary structure elements (Figure 2B and Figure S2B). The fold of the domains is marginally stable towards temperature unfolding, with melting temperatures in the 40-60 °C range (Figure S2A and S2B).

**Figure 2.**
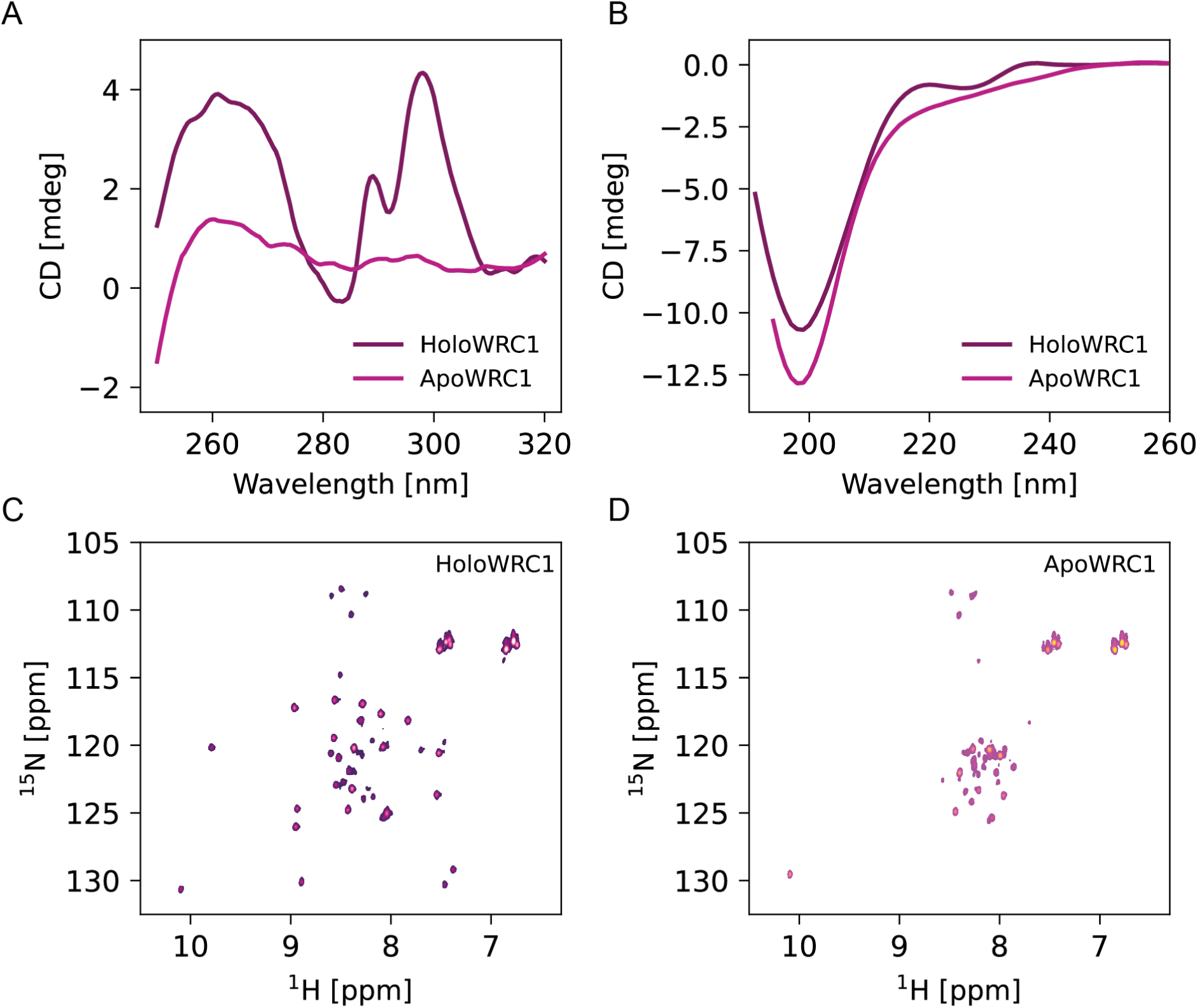
WRC zinc-dependent folding. (A) Near UV CD spectra of WRC1 in the presence (HoloWRC1) and absence (ApoWRC1) of Zn. (B) Far UV CD spectra of Holo and ApoWRC1. 1H-15N HSQC spectra of holo (C) and apo (D) WRC1.

The WRC domain has been proposed as a putative non canonical Zinc Finger domain due to the conservation of three cysteines and one histidine residue. To validate this, we first examined the metal binding properties of the domain using the chelating chromophore PAR^14^. The quantification of the Zn(II) content revealed one equivalent per protein with an affinity in the low nanomolar range (KD = 14.4 nM for WRC7), consistent with the identification of a single zinc-coordination motif (CCCH). We confirmed in this way that WRCs are Zn(II) binding domains (Figure S3).

We then acquired NMR spectra of all four domains. The 1H-15N HSQC spectra show in all cases fewer signals than expected based on the domain’s length and amino acid composition, suggesting the presence of regions with conformational flexibility in an intermediate timescale (Table S5 and Figure S4). Removal of the Zn(II) ion results in loss of the signals’ dispersion, indicating that the metal ion is essential for the domain to acquire its folded conformation (Figure 2, panels C-D and Figure S4). Loss of structure is also reflected in the disappearance of the near-UV CD bands and weak secondary structure elements in the far-UV spectra (∼220 nm) (Figure 2A and 2B, respectively). Therefore, our results confirm that the WRC is CCCH zinc finger domain with one metal binding site which is essential for its native conformation.

### 3.3 The WRC displays a non canonical fold

Unlike canonical CCHH zinc finger domains, which fold into a ββα-motif around a tetrahedrally coordinated Zn²⁺, CCCH zinc fingers adopt non-canonical conformations with limited contributions from defined secondary structure elements ^14,15^. We obtained structural models for the AtGRF1/3/5/7 WRC domains using the AlphaFold3 web server [22]. The resulting structures share a common structural arrangement comprising a β-hairpin stabilized by hydrogen bonds, followed by a loop and a short α-helix. Metal coordination binds these motifs together (Figure 3A and Figure S5). The models show overall little regular secondary structure, in agreement with the far-UV CD spectra.

**Figure 3.**
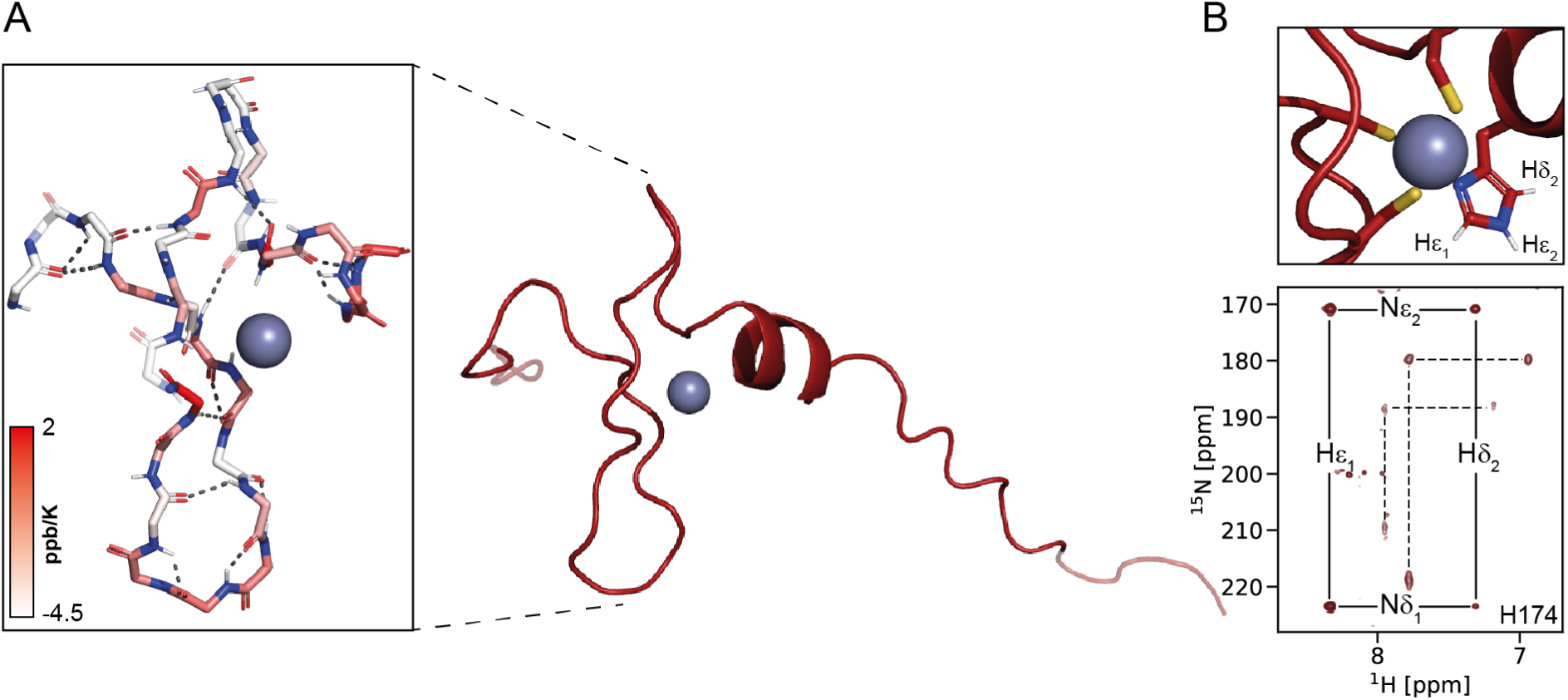
Structural characterization of WRC3. (A) AlphaFold3 model of WRC3. Panel Inset highlights the predicted hydrogen bonding network from the structural mode (dashed lines), with calculated temperature factors (red colormap). The agreement between predicted bonds and experimental NMR data underscores the model’s accuracy. (B) Metal-binding site for which the histidine protonation state is validated by the long-range HSQC spectra displaying characteristic histidine epsilon tautomer side-chain pattern (lower panel).

We proceeded with the NMR assignments for the WRC3 construct to validate the accuracy of the WRC predicted structures (Figure S6). Our NMR analyses indicate that the secondary structure populations derived from WRC3 backbone chemical shifts align well with the predicted secondary structure elements (Figure S7). Furthermore, hydrogen bond interactions, assessed through ^1^H_N_ temperature coefficients, closely match the predicted H-bond networks, reinforcing the model’s accuracy (Figure 3A, inset, and Figure S7). Perturbation of the secondary structure with trifluoroethanol (TFE) leads to linear displacements of the ^1^H_N_ chemical shifts. The changes in the slope of these displacements correlate well with the boundaries of secondary structure motifs in the model, further validating the predicted locations of these elements (Figure S7). The WRC3 HetNOE profile along with R1 and R2 NMR relaxation measurements confirm the presence of unstructured N- and C-terminal segments outside the predicted folded regions (Figure S8).

Additionally, we confirmed the orientation of the folded structural elements and their relative spatial distribution via H_N_-N residual dipolar coupling (RDC) and NOESY-HSQC analyses. RDC measurements of amide protons in WRC3 within an aligned medium indicate that the relative orientation of the α-helix and β-hairpin matches the model’s predicted orientation (Figure S7). The cross-peaks observed in NOESY-HSQC spectra agree with those expected from the model, further supporting the spatial distribution and predicted inter-residue distances for several WRC amino acids (Figure S9).

Finally, we assigned side-chain resonances of the Zn-bound histidine to analyze its tautomeric state (Figure S10). The signal pattern observed in long-range 1H-15N HSQC spectra for all constructs reveals a histidine in a HNε tautomeric form, indicating that the metal ion is coordinated via Nδ (Figure 3B, Figure S11) ^36^. On top of that, we also observed NOEs between the aromatic proton resonances of the conserved tryptophan and the coordinating histidine (Figure S10). All of these results agree with the AlphaFold3 model, suggesting that the modeled structure efficiently predicts the WRC folding at the side-chain level, at least for the coordinating histidine.

In contrast to other canonical Zn fingers, WRC domains lack a hydrophobic core (Figure S5D). Regarding the conserved WRC motif, the cysteine functions as one of the Zn(II) ligands, while neither the tryptophan nor the arginine side chains appear to play a direct role in specific interactions within the domain. The presence of a basic residue preceding the metal-coordinating cysteine has been linked to a reduction in the pKa value of the thiol, thereby enhancing the metal site’s affinity [23,24]. On the other hand, the tryptophan residue may be involved in nucleic acid interactions, as observed in other similar domains.

### 3.4 The WRC motif is involved in DNA recognition

The WRC domains of GRFs ^6,7^ and other proteins ^17^ were found to drive interactions with dsDNA segments in promoter regions. To explore the mechanisms underlying DNA recognition and binding by WRCs, we conducted *in vitro* biophysical studies on the WRC domains of *A. thaliana* GRFs.

Our analysis focused initially on WRC7, as its DNA target sequence is more thoroughly studied than those of other GRF WRCs ^7^. To assess WRC binding, we evaluated its interaction with previously reported specific and non-specific DNA sequences using tryptophan fluorescence quenching ^37^. We observed that all DNA fragments quenched the intrinsic fluorescence of WRC, regardless of their specificity (Figure 4A). However, the dsDNA containing the cis-targeting sequence induced a complete fluorescence quenching, whereas the non-specific sequence resulted in only partial quenching. Complete quenching was also observed for a minimal sequence harboring the specific DNA binding motif (Figure S12). This behaviour suggests a preference for specific DNA sequences and points toward a possible role for the conserved Trp in DNA binding. Moreover, CD difference spectra of WRC in presence of the minimal dsDNA target sequence also show significant changes (Figure 4B). The near UV region shows an intense unstructured difference band, suggesting that WRC binding causes significant alterations in the conformation of bound DNA. The far UV region shows a ca. 7 nm red shift in the minimum, hinting at a rearrangement of the WRC backbone. The titration curves obtained by both methodologies allowed us to estimate an occupation of ca. 7 bp for WRC7, consistent with the length of the reported GRF7 recognition sequence.

**Figure 4.**
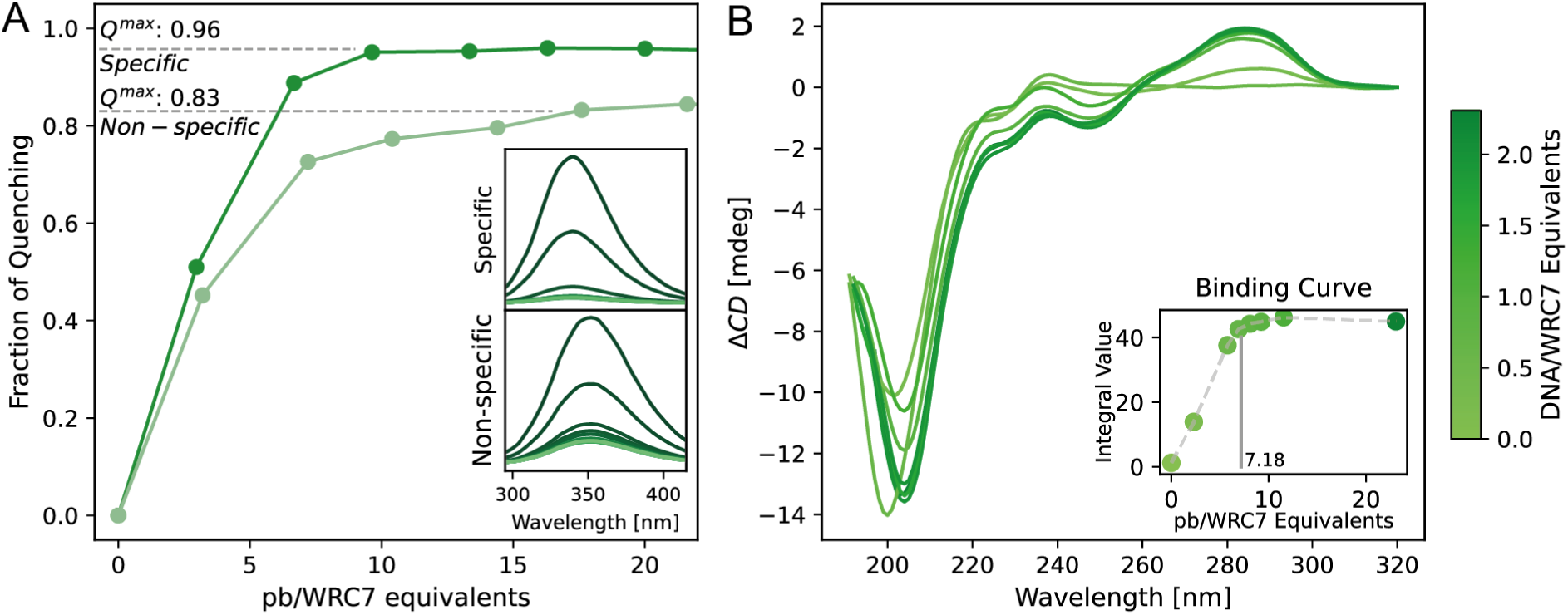
WRC7 DNA-binding assays. (A) Tryptophan fluorescence quenching of specific (dark green) and non-specific (light green) DNA sequences at different pb/WRC7 equivalents. Maximal quenching for each probe is indicated by dashed lines. Insets show the fluorescence emission spectra for each pb/WRC7 ratio, with darker colors representing lower ratios. (B) Circular dichroism (CD) difference spectra of WRC7 titrated with DNA. The inset displays the binding curve derived from the integral CD signal, indicating an occupancy of 7.18 pb per WRC molecule.

We then aimed to identify the residues involved in WRC-DNA interaction using NMR spectroscopy. We studied the WRC1 domain due to its higher stability and the larger number of signals observed in the HSQC spectra compared to WRC7 (Figure S2C and S4, and Table S5). We designed minimal specific and non-specific binding probes based on previously reported GRF1 cis-targeting sequences and confirmed their binding to WRC1 by CD spectroscopy (Figure S13). To map the residues involved in WRC-DNA interaction, we acquired NMR spectra of WRC1 in the presence of either target DNA or control DNA. Surprisingly, while the complex with target DNA gives a well resolved spectrum, the complex with control DNA yields a spectrum with fewer signals (Figure S13A and Figure S13B, respectively). These results suggest that, while WRC1 binds both specific and non-specific DNA, it only forms a well-defined complex in the case of the specific DNA but an undefined complex in the case of the non-specific DNA. This finding aligns with the differences in specificity observed for WRC7 probes and accounts for the higher stabilization of the WRC-DNA complex in the presence of specific DNA sequences.

We calculated the chemical shift perturbation of assigned signals in the presence of the specific minimal sequence (Figure 5A and Figure S13A). As shown in Figure 5B, residues exhibiting significant chemical shift perturbations cluster into the same region within the WRC folded structure. Residues G200, R201, K209, W210, R211, C212 and R225 form the core DNA recognition interface. In addition, the signal corresponding to residue E198 which lies at the end of the disordered N-terminal region and in close proximity to G200 and R201 disappears upon DNA addition, also suggesting a role in DNA binding. Zinc removal was found to impair dsDNA binding (Figure S13), consistent with the assembly of a DNA interaction surface that integrates the two distant arginine residues found in positions 201 and 225 and the conserved WRC motif. However, due to the incomplete WRC assignment we cannot rule out the participation of other residues in the WRC-DNA interaction. Notably, E198, G200, K209, and R225 are key residues that define the unique amino acid signature of GRF-type WRCs, as revealed by our bioinformatic analysis (Figure 1). This signature may underlie the sequence specificity of WRCs within this family of transcription factors. Our findings highlight the role of the conserved WRC motif in DNA binding and suggest that zinc binding is not only essential for structural integrity but also for the proper alignment of residues critical for DNA recognition.

**Figure 5.**
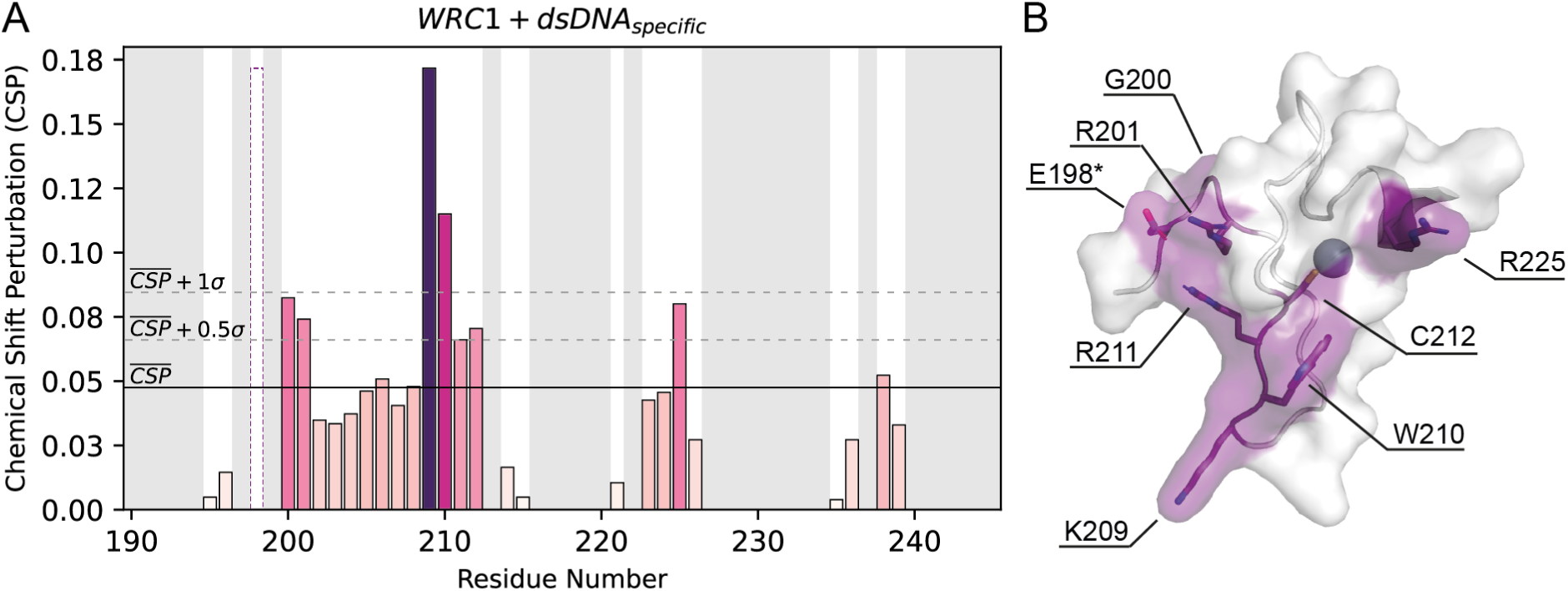
DNA-binding interaction of WRC1 with the specific minimal probe. (A) Chemical shift perturbation (CSP) analysis of WRC1 residues upon titration with specific dsDNA. Bars are colored proportionally to the CSP values, with darker colors indicating larger perturbations. The dashed bar corresponds to residue E198, which disappears upon DNA binding, indicating a significant conformational change or exchange broadening. Thresholds corresponding to the mean (CSP), mean + 0.5σ, and mean + 1σ are indicated by solid and dashed lines. Gray-shaded areas represent non-assigned residues. (B) Mapping of the perturbed residues with a CSP higher than mean + 0.5σ onto the folded region of the validated WRC3 structure. Key residues involved in DNA binding are labeled. Residues in both panels are numbered based on the GRF1 primary sequence.

## 4 Discussion

Growth-regulating factors (GRFs) play integral roles in diverse biological processes that shape plant growth, development, and physiological responses ^12,38^. The critical functions of this transcription factor family and their potential biotechnological applications have sparked considerable interest in elucidating their molecular mechanisms. Numerous studies have focused on identifying the DNA cis-regulatory sequences targeted by GRFs across various plant species ^7,9,13,35^. As transcription factors, GRFs exhibit a modular domain architecture comprising three distinct regions: the QLQ domain for protein-protein interactions, the WRC domain for DNA binding, and a C-terminal transactivation region. Since their discovery, the WRC domain has been hypothesized to function as a zinc finger (ZF) due to its similarity to the CCCH motif of the Hordeum Repressor of Transcription (HRT) DNA-binding domain ^1,10^. However, the mechanisms by which the WRC domain enables GRFs to recognize and bind their DNA targets remain poorly understood. In this study, we provide a comprehensive characterization of the structural and functional properties that distinguish the WRC domains within the GRF family.

Our bioinformatics and experimental analyses reveal a unique amino acid signature within GRF-associated WRC domains, which appears essential for their DNA recognition specificity. Furthermore, singular value decomposition (SVD) analysis underscores the distinctive nature of these domains by clustering GRF WRCs into a discrete group, reinforcing the notion that their intrinsic features are tailored for specific biological roles.

ZF transcription factors represent a versatile class of DNA-binding proteins characterized by zinc-coordinating motifs that stabilize their structure and enable DNA interaction, and account for one of the largest transcription factor families in plants ^39,40^. Whereas many characterized ZF can be grouped based on a limited set of canonical structures, these domains present an extremely large variability in terms of global conformation. The structural validation we present here confirms that WRCs are CCCH ZF domains that adopt a non-canonical fold, with the metal ion playing a crucial role in stabilizing their conformation. We found that WRCs lack a distinct hydrophobic core, making zinc coordination and hydrogen bonding the primary forces driving their folding into a specific tertiary structure. The low quality of their NMR spectra, with signals absent for several residues in HN-N and triple resonance spectra, indicates that even the core region is flexible in a µs-ms timescale. This flexibility may be important for binding a variety of targets. In fact our experiments suggest that WRCs can bind both specific and non-specific dsDNA. The discrimination between both most probably needs local conformational rearrangements of the backbone structure that are explored in the free protein.

WRC domains appear to be necessary for binding of GRFs to DNA, but not sufficient to confer specificity. Transcription factors that regulate developmental processes, such as GRFs, must be tightly controlled and limited to particular cell types and precise developmental stages. In the present work we study the domain isolated from its native context. One could speculate that the missing information required for sequence specificity may reside in interactions with cellular partners, adding a further control layer to the GRFs regulatory network.

Within the defining triad of residues WRC, the cysteine is one of the Zn(II) ligands and thus essential for the stability of the fold, but neither the arginine nor the tryptophan make significant contacts with the rest of the protein. Instead their sidechains are both exposed and appear to be essential for nucleic acid binding, as reported by the NMR spectra of the complex. In RanBP2-type ZnFs a tryptophan and two arginines dictate the recognition of specific nucleotides in single stranded RNAs, with the tryptophan stacking between two bases ^41,42^. Although this binding mechanism is not feasible for double standard DNA given its conformational restraints, our results along with the high conservation of this residue within WRCs hint towards a pivotal role for this aromatic amino acid in DNA binding. Further studies will be essential to delineate the structural function of the indole sidechain in the formation of GRFs:DNA complexes in atomic detail.

GRFs are among the few known CCCH ZFs that bind directly to DNA to regulate gene expression ^43,44^. This contrasts with the majority of CCCH zinc finger proteins, which are primarily associated with RNA metabolism, including RNA cleavage, degradation, polyadenylation, and other post-transcriptional processes ^45–49^. Moreover, while HRT and other CCCH proteins have multiple ZF domain copies, most GRFs only harbor one WRC domain. To our knowledge, GRFs function has only been linked to DNA association, however the unique structural and biochemical features behind GRFs’ architecture make this system appealing for the exploration of new biological roles. We believe that the present study of WRC domains from GRF transcription factors may inspire the development of new hypotheses and guide future experiments, contributing to a deeper understanding of this important family of plant transcription factors.

## 5 Author Contributions

**Franco A. Biglione:** Conceptualization, data curation, experimental procedures, visualization, formal analysis, writing initial draft, review and editing. **Javier F. Palatnik:** Conceptualization, writing review and editing. **Nahuel D. González Schain:** Conceptualization, funding acquisition, project administration, supervision, writing review and editing. **Rodolfo M. Rasia:** Conceptualization, formal analysis, funding acquisition, project administration, supervision, writing initial draft, review and editing.

## Acknowledgments

F.A.B. is a Fellow at CONICET. J.F.P., N.D.G.S and R.M.R. are Career Researchers at CONICET. We are grateful to Silvana Sut for her excellent assistance with labware and media preparation, as well as Andrea Coscia and Alejandro Gago for maintenance of the NMR facility. We also thank Daniela Liebsch for insightful discussions and valuable input throughout the development of this work.

---

**Table S1.** Protein domain architectures of WRC-containing protein sequences and their distribution in each cluster. Domain architectures were retrieved from UNIPROT domain names describing each protein entry.

| *Domain architecture* | *Cluster gray* | *Cluster gold* | *Cluster blue* |
| --- | --- | --- | --- |
| QLQ_WRC | 0 | 346 | 0 |
| WRC | 119 | 81 | 265 |
| WRC_RING-type_JmjC | 0 | 0 | 147 |
| QLQ_WRC_WRC | 0 | 101 | 1 |
| WRC_WRC | 0 | 46 | 20 |
| WRC_JmjC | 0 | 0 | 36 |
| WRC_RING-type | 0 | 0 | 27 |
| WRC_WRC_WRC | 0 | 1 | 14 |
| WRC_WRC_WRC_WRC | 0 | 0 | 6 |
| WRC_WRC_WRC_WRC_WRC | 0 | 0 | 5 |
| WRC_WRC_RING-type | 0 | 0 | 3 |
| WRC_WRC_WRC_RING-type | 0 | 0 | 3 |
| WRC_WRC_WRC_WRC_JmjC | 0 | 0 | 3 |
| WRC_WRC_WRC_WRC_WRC_WRC | 0 | 0 | 3 |
| QLQ_WRC_WRC_WRC | 0 | 2 | 1 |
| WRC_WRC_WRC_WRC_RING-type_JmjC | 0 | 0 | 2 |
| CCHC-type_QLQ_WRC | 0 | 1 | 0 |
| DEK-C_WRC | 0 | 1 | 0 |
| GATA-type_GATA-type_QLQ_WRC | 0 | 1 | 0 |
| QLQ_WRC_QLQ_WRC | 0 | 1 | 0 |
| QLQ_WRC_SHSP | 0 | 1 | 0 |
| C2H2-type_WRC | 0 | 0 | 1 |
| F-box_WRC | 0 | 0 | 1 |
| GATA-type_WRC | 0 | 0 | 1 |
| JmjN_JmjC_WRC | 0 | 0 | 1 |
| WRC_Myb-like | 0 | 0 | 1 |
| WRC_RING-type_JmjC_Peptidase A1 | 0 | 0 | 1 |
| WRC_SET_Post-SET | 0 | 0 | 1 |

**Table S2.** Distribution of protein groups across WRC clusters. GRF-type and JmJ-C type group WRC-containing proteins with domain architectures harboring at least one QLQ domain and JmjC domains (table S1), respectively. Protein for which the only domain annotation corresponds to WRC domains are names according to the number of WRCs present (WRC_1_ and WRC_n_ for only one or more than one WRC domains, respectively). Proteins with annotated domains different from QLQ or JmjC are named as Other.

| *Protein groups* | *Cluster gray* | *Cluster gold* | *Cluster blue* |
| --- | --- | --- | --- |
| GRF-type | 0 | 453 | 2 |
| JmjC-type | 0 | 0 | 190 |
| WRC<sub>1</sub> | 119 | 81 | 38 |
| WRC<sub>n</sub> | 0 | 47 | 265 |
| Other | 0 | 1 | 48 |

**Table S3.**
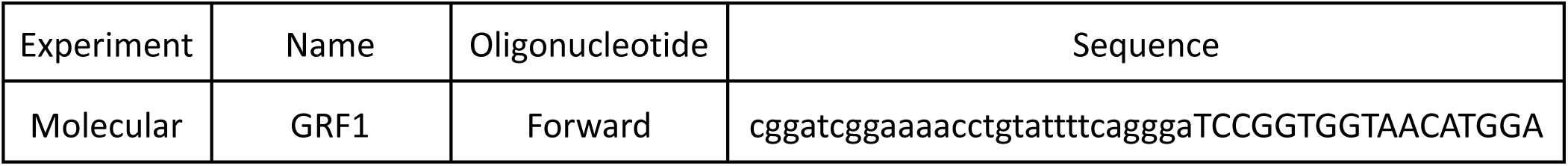
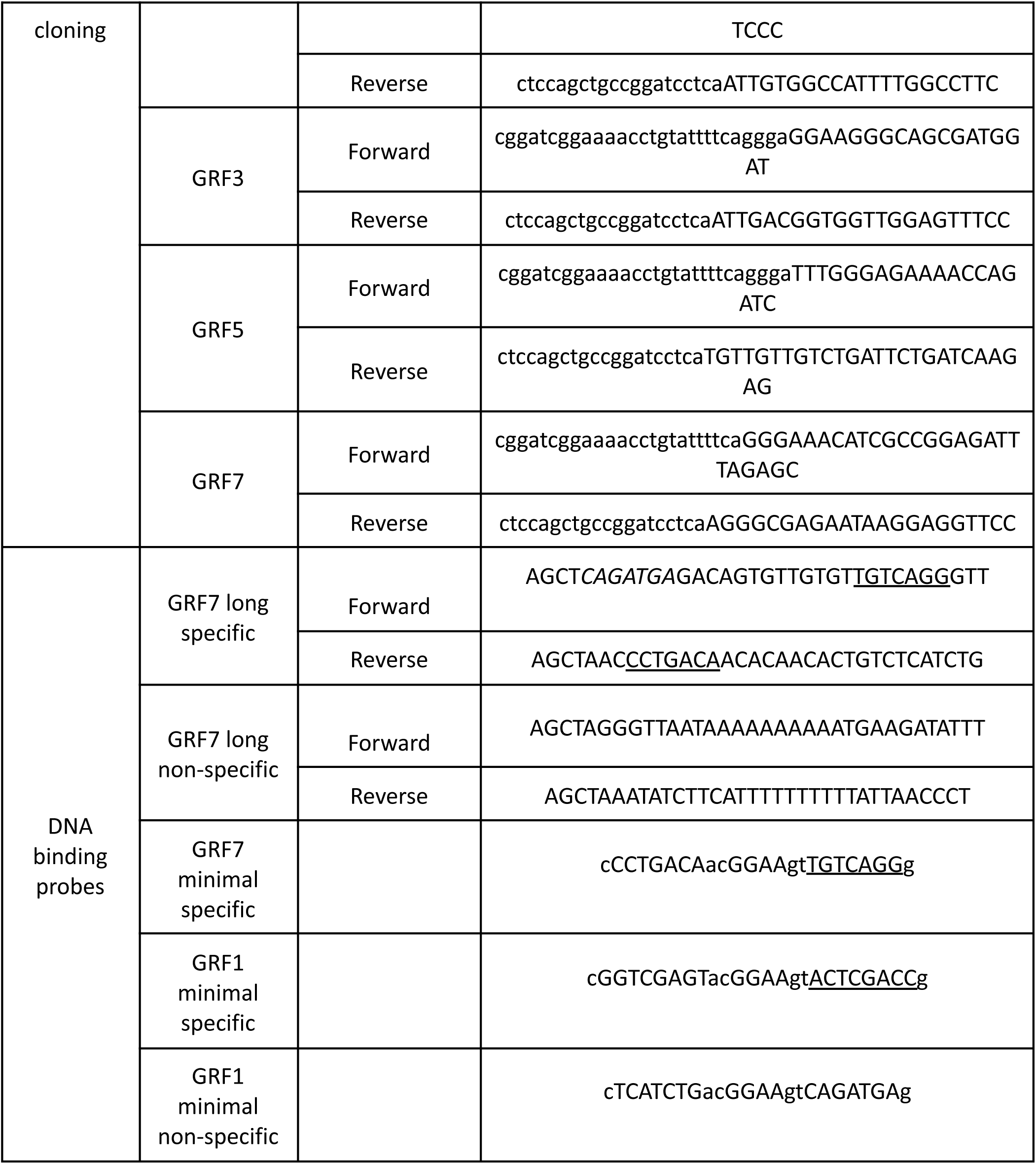
Oligonucleotide sequences used in this study. Sequences used for restriction-free (RF) cloning of WRC sequences are listed under the “Molecular Cloning” row, with regions complementary to the destination vector shown in lowercase. Oligonucleotides used in DNA binding assays are listed under the “DNA Binding Probes” section, with reported GRF cis-targeting sites underlined. For minimal sequences, lowercases denote opening and closing base pairs within the stem region. Nucleotides within the GRF7 long specific oligonucleotide formatted in italic correspond to a region shown not to interact with GRF7 in the source publication, and was used as GRF1 minimal non-specific probe for NMR studies.

**Table S4.**
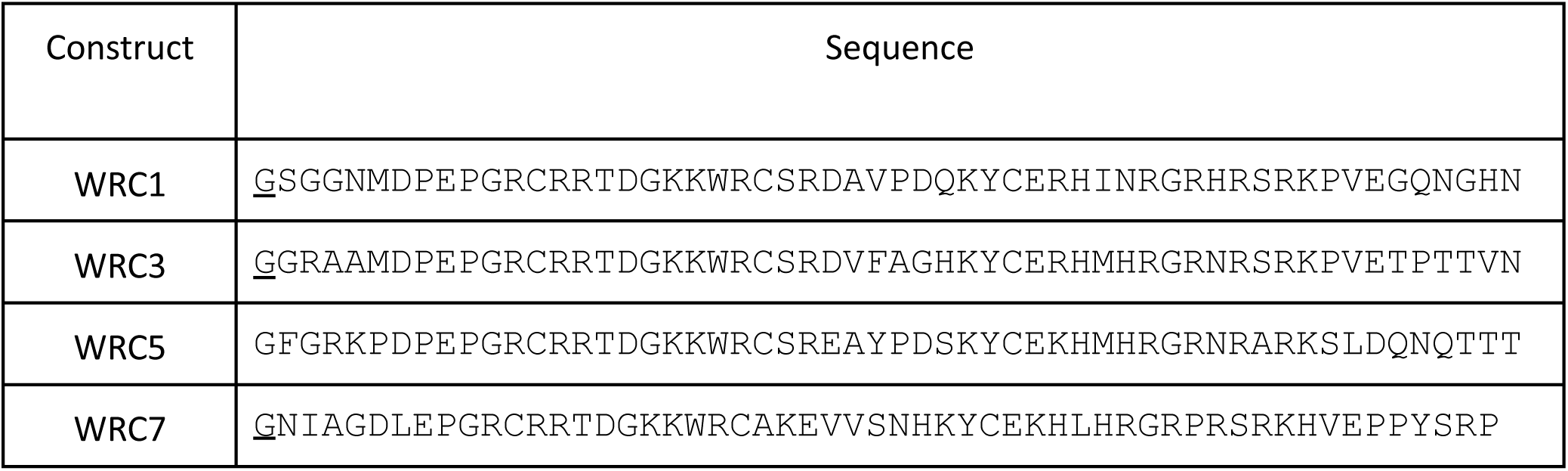
Sequence of purified WRC variants. Underlined residues correspond to remaining glycines after cleavage at the TEV recognition site (ENLYQF/G) not present in the original GRF sequences.

**Table S5.** Number of backbone ^1^H-^15^N cross peaks expected vs observed in HSQC spectra.

| Construct | Zn(II) containing protein |  |
| --- | --- | --- |
|  | Expected | Observed |
| WRC1 | 51 | 43 |
| WRC3 | 51 | 53 |
| WRC5 | 51 | 46 |
| WRC7 | 49 | 39 |

## 7 Supplementary information

**Figure S1.**
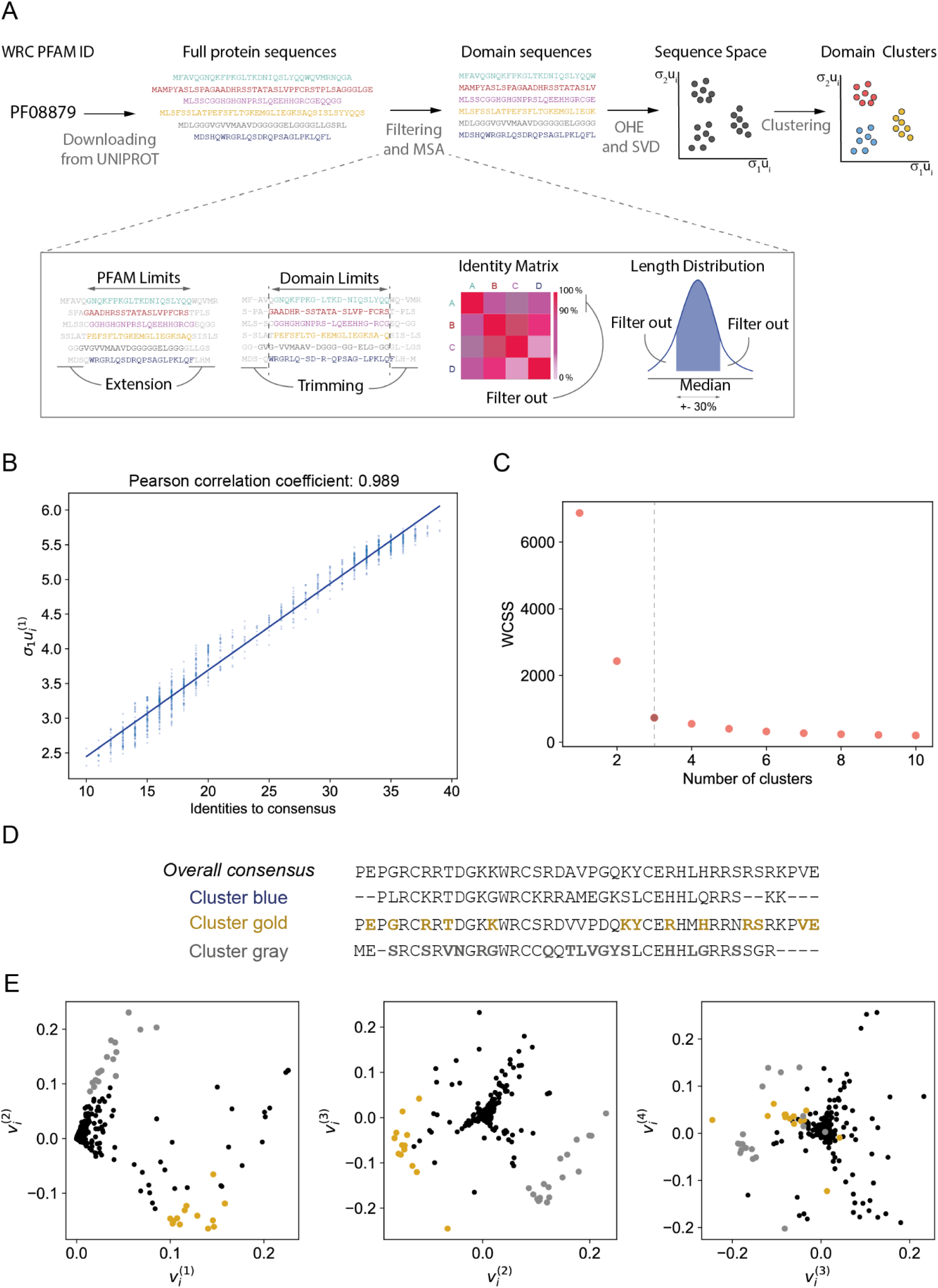
Singular value decomposition of WRC sequences (A) Graphic summary of the bioinformatic pipeline performed in the analysis. (B) Correlation between consensus identity of MSA sequences and σ^1^u_i_^(1)^ values. The x-axis gives the average number of identities of each sequence with the consensus sequence determined from the entire MSA. The solid blue line indicates a linear least-squares fit, with the Pearson correlation coefficient displayed in the panel title. (C) Elbow plot for WRC, showing clustering based on the first three σ_k_u_i_^(k)^ values. The within-cluster sum of squares (WCSS) decreases sharply until k=3, indicating an optimal number of three clusters. (D) Overall and cluster-specific consensus sequences obtained from SVD analysis. Residues of each cluster with a preferential in-cluster vs out-of-cluster enrichment of at least 60% are highlighted by bold colored letters. (E) Residue eigenvectors of the SVD of WRC sequences. The residues that show the higher contribution to cluster separation are highlighted by colored markers. Sequence clusters and cluster-enriched residues are in the same directions in SVD space.

**Figure S2.**
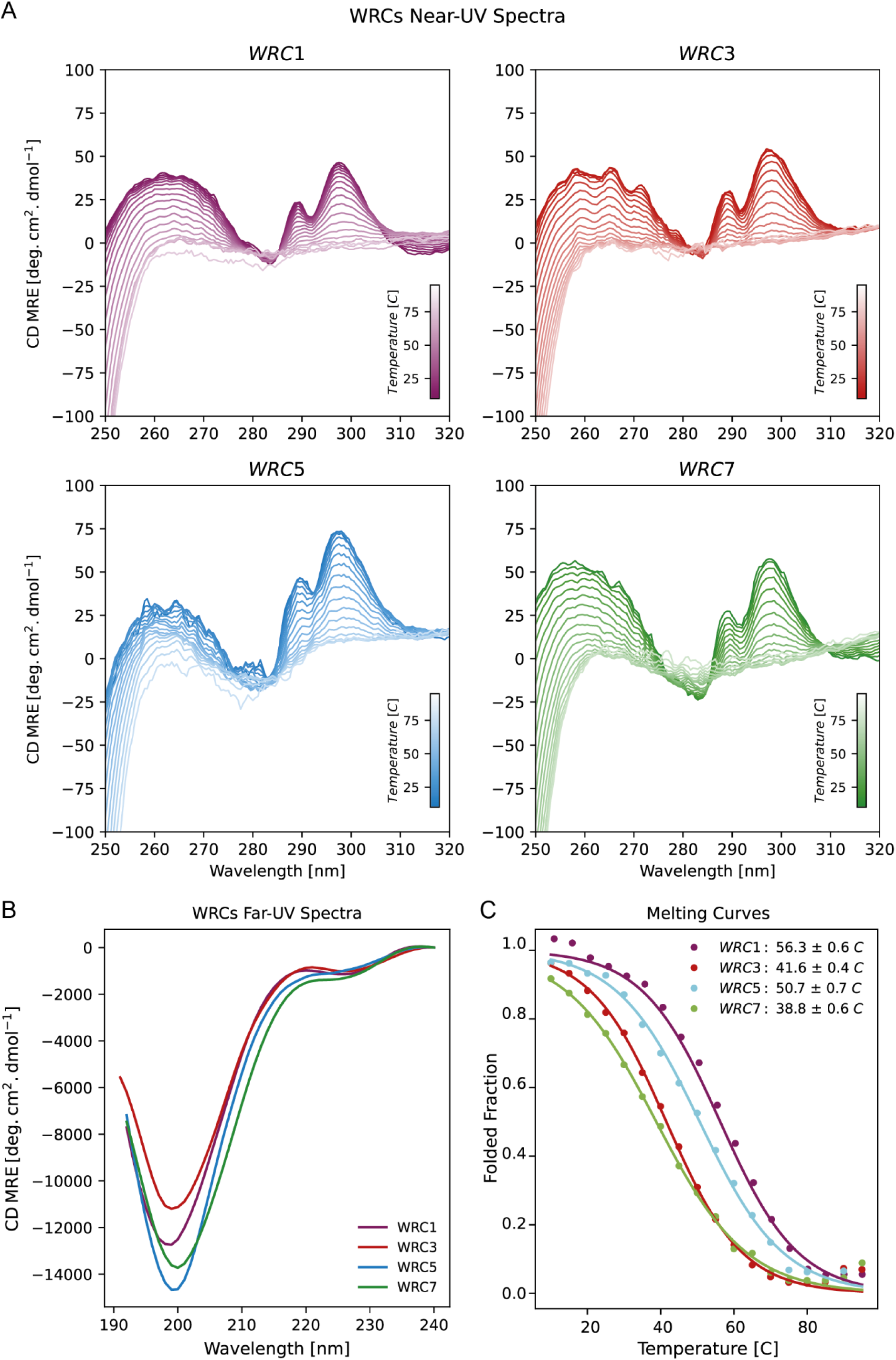
Overall folding of WRC domains by CD spectroscopy. Near-UV (A) and far-UV (B) spectra of WRCs. Melting curves derived from near-UV spectra acquired at increasing temperatures were fit to a two-state unfolding model (C).

**Figure S3.**
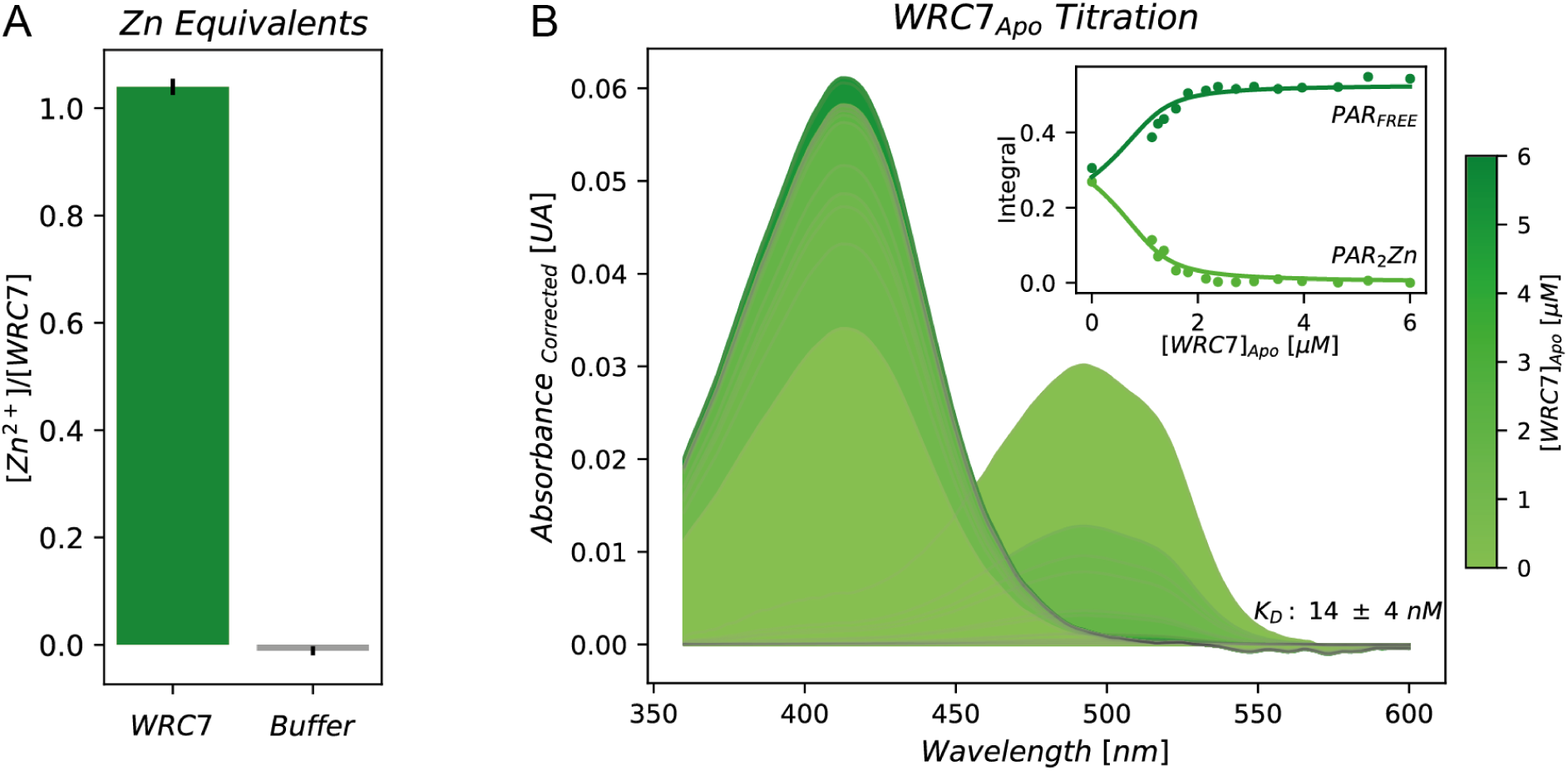
Zn^2+^ binding stoichiometry (A) and affinity (B) of WRC7 measured by competition with PAR. Integrals of a 10 nm window from absorption maxima of deconvoluted PAR spectra under increasing WRC concentrations were fitted to a one-site binding model (Panel B, inset).

**Figure S4.**
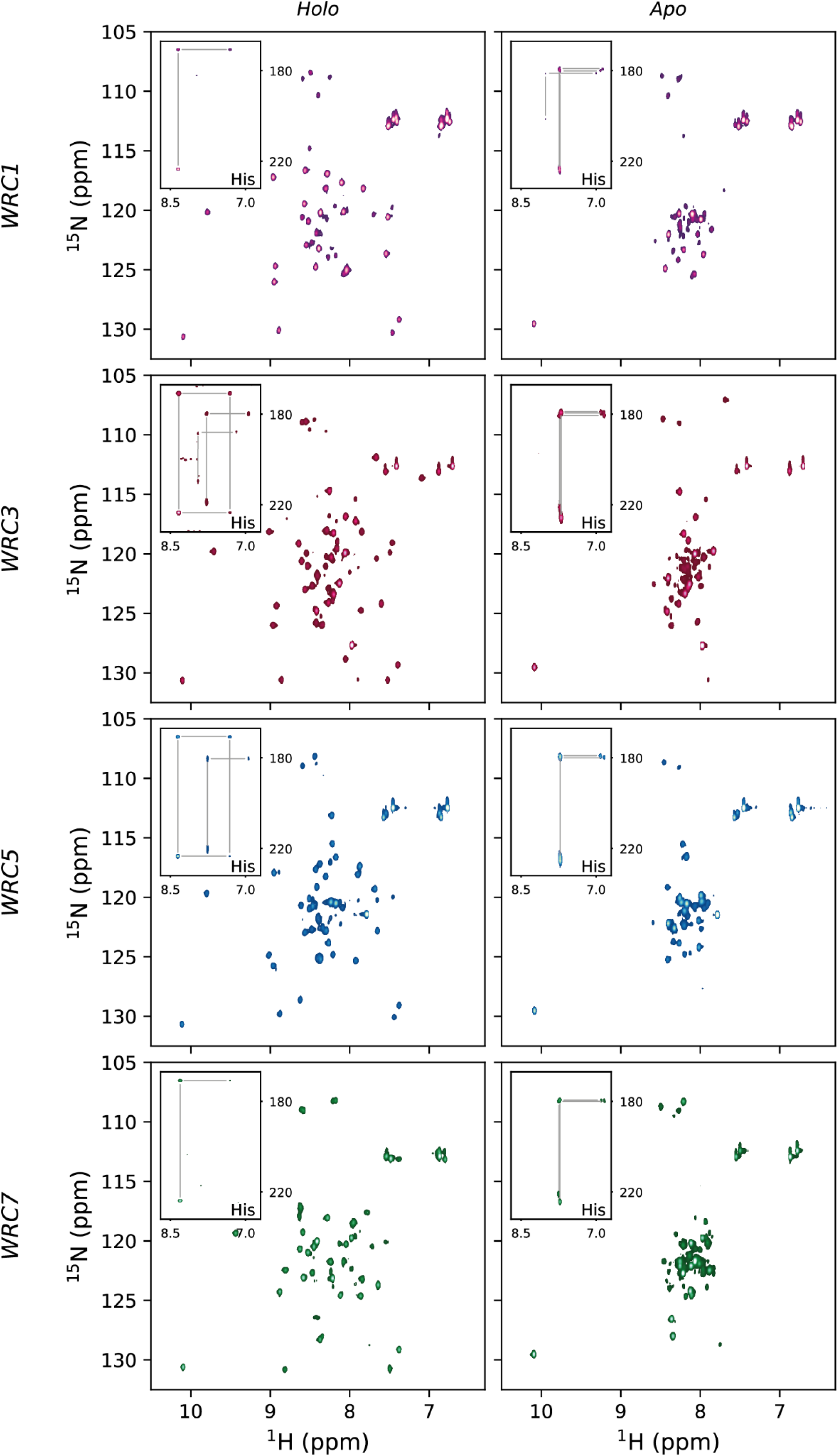
Zn(II) loss induces unfolding, as evidenced by ^1^H-^15^N HSQC spectra. The signals corresponding to the histidine side-chain resonances (insets), which display a conserved pattern across all isoforms, are highly sensitive to the metalation state of the WRCs. This sensitivity highlights the involvement of at least one histidine side chain in coordinating Zn(II), thereby playing a critical role in maintaining the overall structure of the domain.

**Figure S5.**
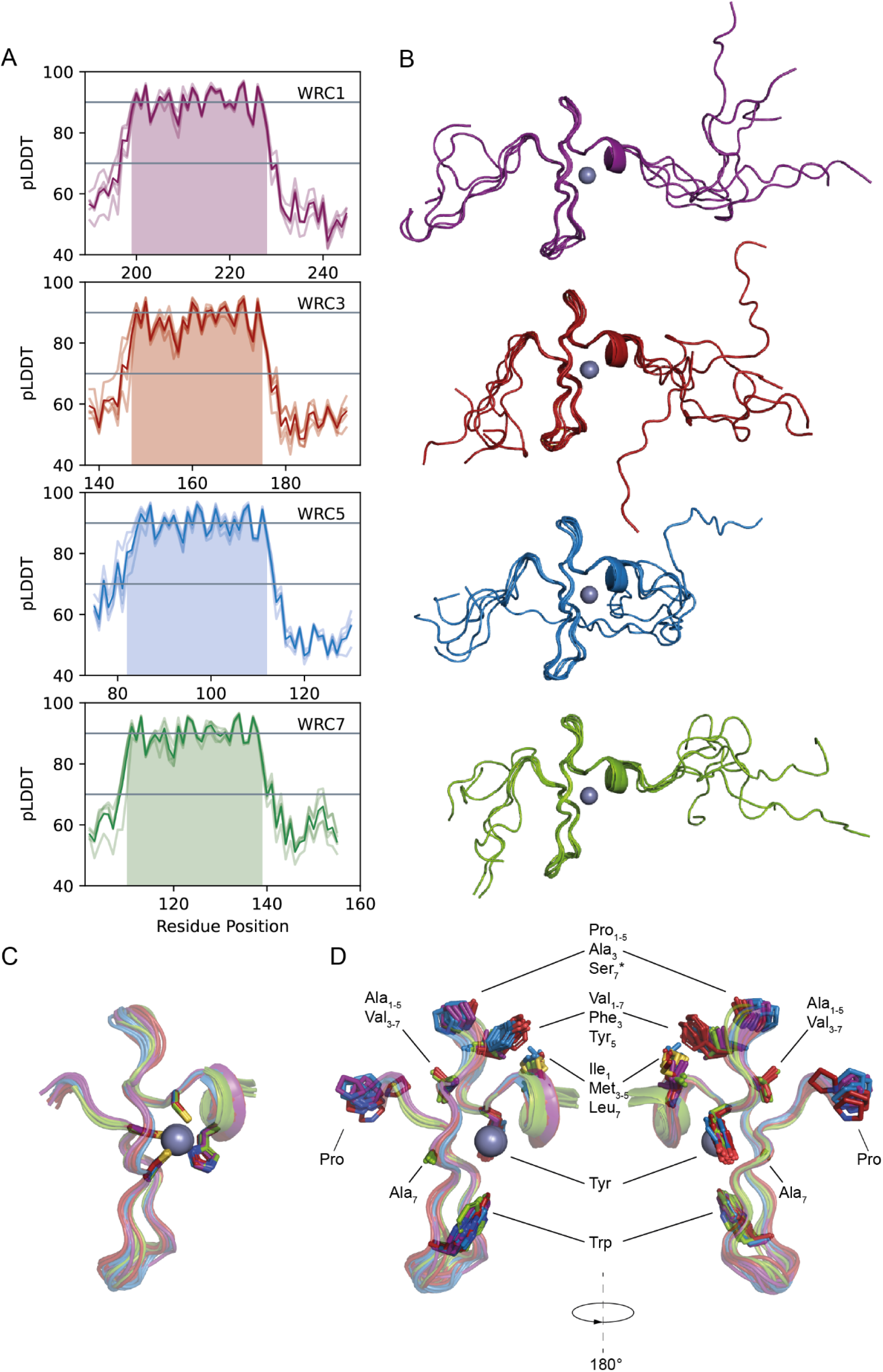
Structural models calculated with AlphaFold3. (A) pLDDT profiles of the five ranked models for each calculated WRC structure (shadowed lines). Solid lines represent the overall mean pLDDT per residue, position numbering corresponds to GRF1, GRF3, GRF5 and GRF7. Very high (pLDDT > 90) and moderate (pLDDT > 70) confidence ranks are indicated by gray lines. Colored areas highlight regions where pLDDT exceeds a threshold value of 80. (B) AF3 structural models for each WRC aligned in regions spanning confident pLDDT values (pLDDT>80) . Zinc ion is displayed as a green sphere. (C) Side chain structural assessment of the CCCH metal coordination site. (D) Distribution of non-polar residues within the folded regions of the WRC models. Subindices denote the WRC isoform containing the amino acidic variant at each position. Polar residue variants at mainly non-polar positions are indicated by an asterisk.

**Figure S6.**
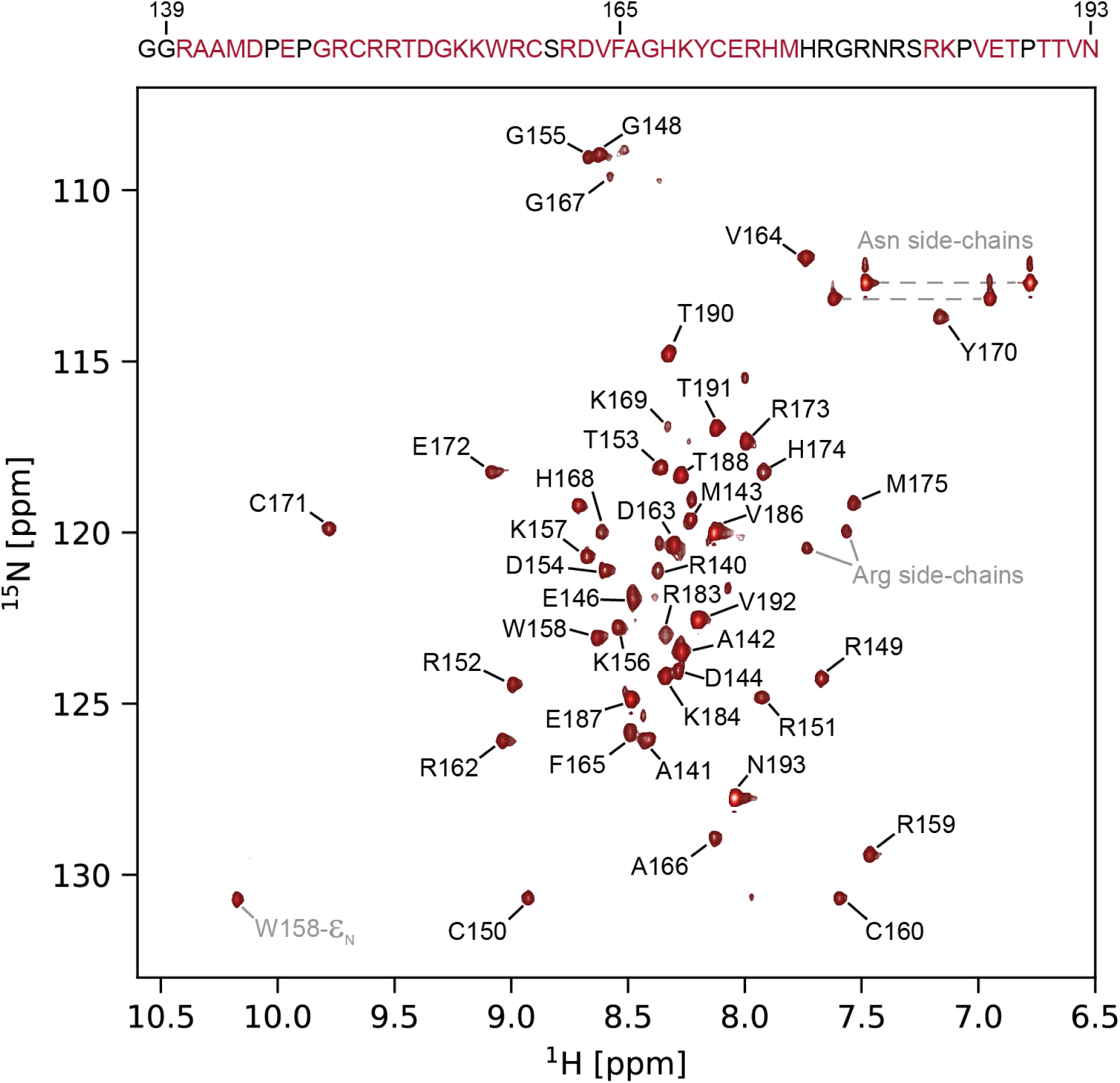
NMR backbone assignment of WRC3. ^1^H-^15^N HSQC spectra of WRC3 at 298 K. Backbone and sidechain resonances are indicated by black and gray labels, respectively. Arginine side-chains were identified by signal folding in ^15^N-^1^H SOFAST-HMQC spectra.

**Figure S7.**
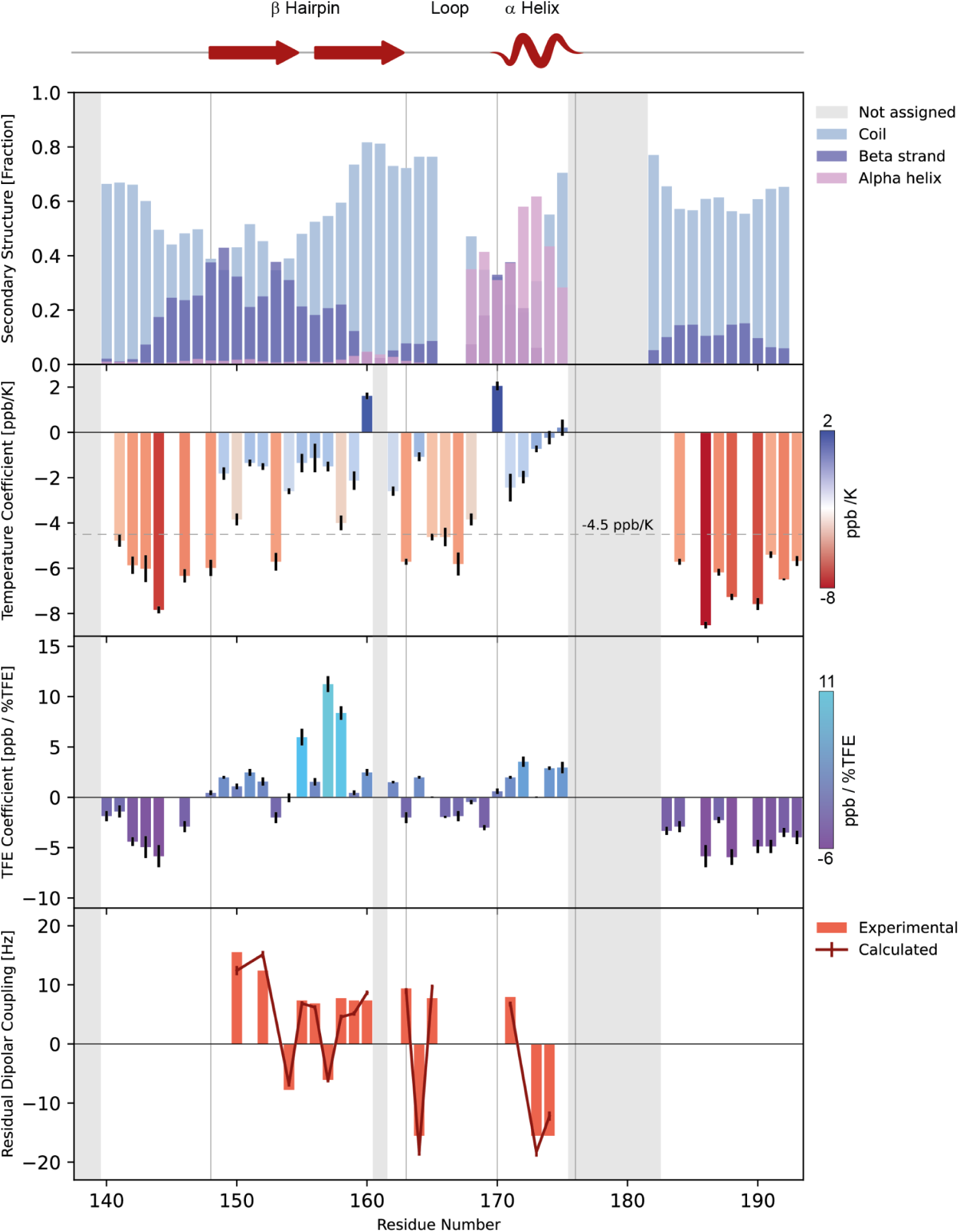
Structural validation of AF3 WRC3 structure. First row: Global analysis of the secondary structure propensities considering all backbone chemical shifts using d2D software. Second row: Temperature coefficients calculated from amide chemical shifts (second row). Residues above the threshold ( −4.5 ppb/K, dashed line) likely participate in hydrogen bond formation. Third row: Secondary structure mapping through TFE perturbation coefficients. Fourth row: Experimental and calculated H_N_-N residual dipolar couplings (RDCs) of the folded WRC regions. Gray-shaded areas represent non-assigned residues. Vertical lines denote the limits of the secondary structure elements depicted in the AF3 model (top illustration). Residues are numbered according to GRF3 primary sequence.

**Figure S8.**
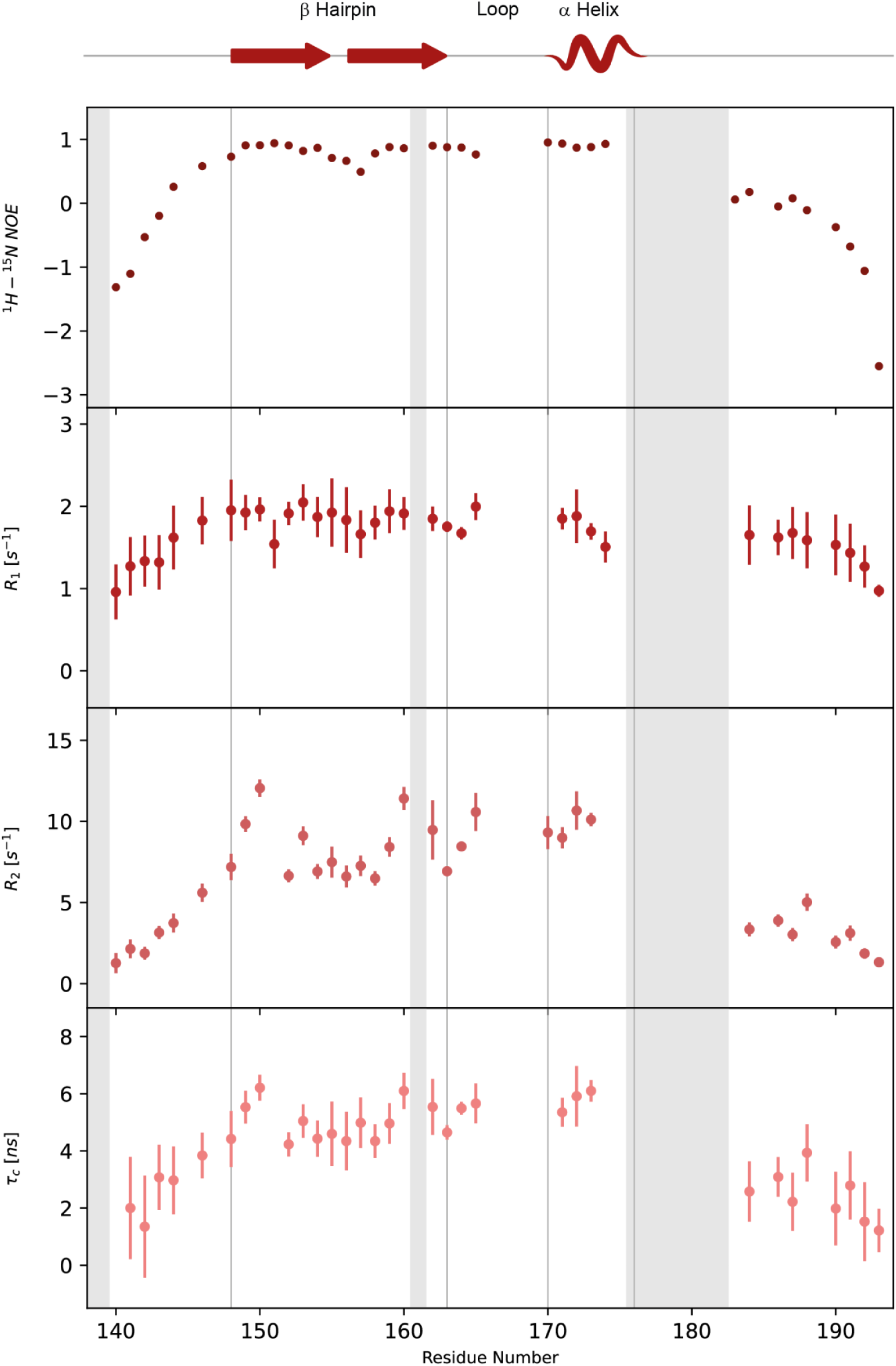
NMR relaxation dynamics of WRC3 assessed through ^1^H-^15^N HetNOE, R1 and R2 data. Correlation time (tc) was estimated from the R2/R1 ratio. Comprehensive interpretation of the fourth parameters and determination of flexible regions allowed the identification of unstructured and folded segments within the protein. Gray-shaded areas represent non-assigned residues. Vertical lines denote the limits of the secondary structure elements depicted in the AF3 model (top illustration). Residues are numbered according to GRF3 primary sequence.

**Figure S9.**
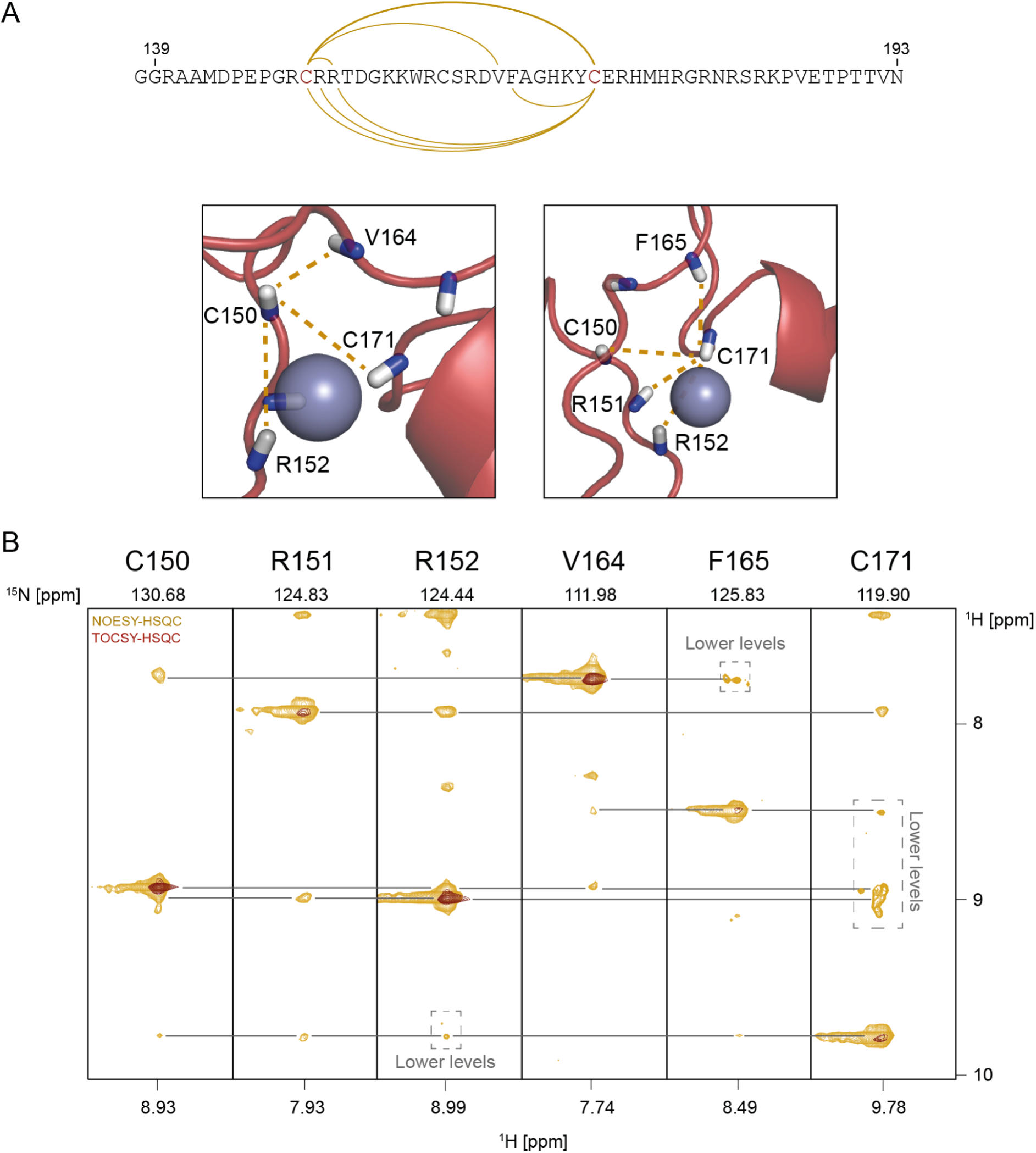
Validation of structural constraints in the WRC3 model. (A) Structural representation of the AF3 coordination site, highlighting residues C150 and C171, which are spaced in the primary sequence. NOE interactions between residues and cysteines (gold lines) are mapped to confirm spatial proximity in the model. (B) TOCSY-HSQC and NOESY-HSQC spectra, showing cross-peaks that support the spatial distribution and predicted inter-residue distances in the WRC3 domain. Residues are numbered according to GRF3 primary sequence.

**Figure S10.**
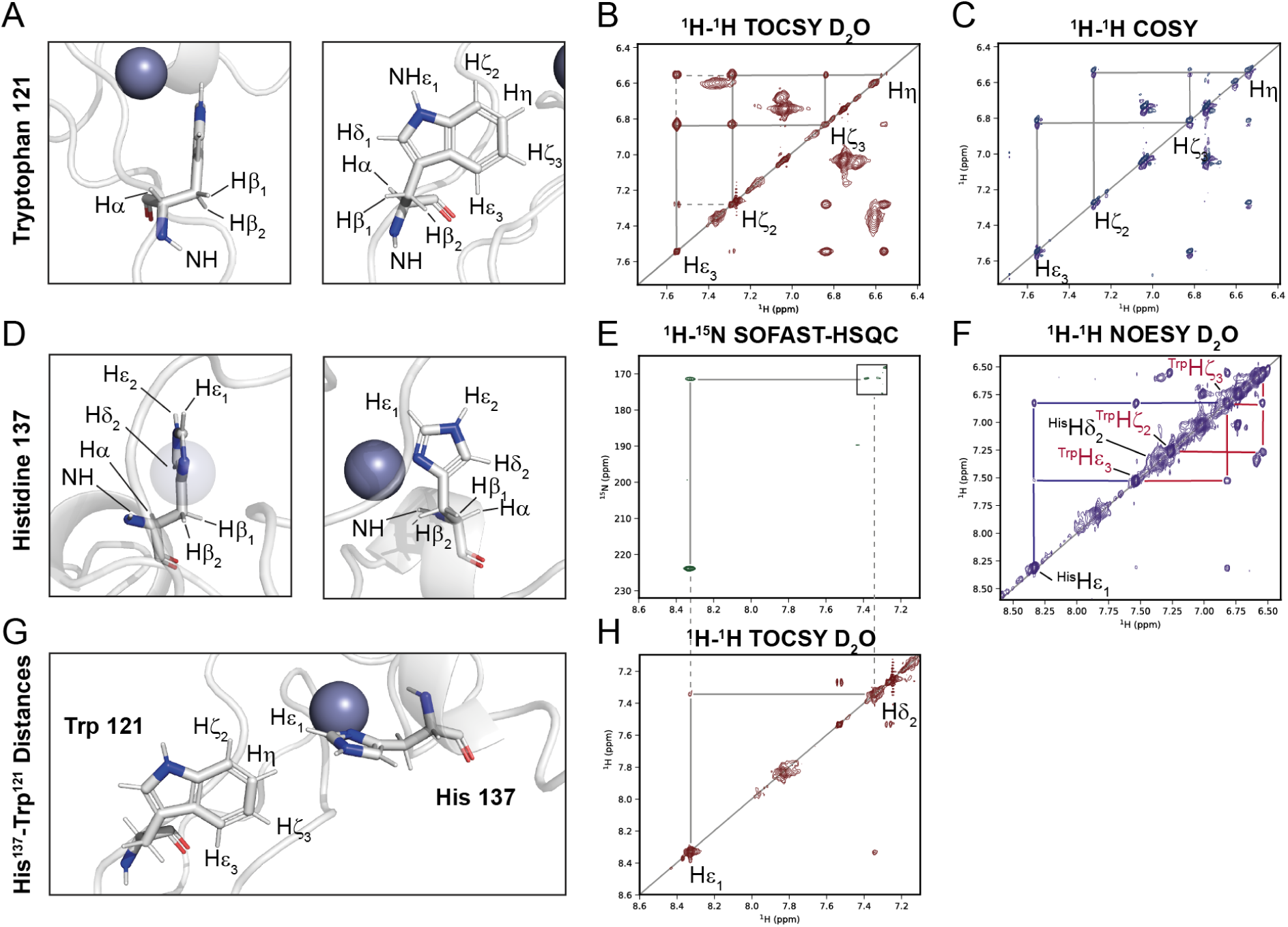
Assignment and validation of histidine and tryptophan side-chain resonances in the WRC7 model. (A, D, G) Structural model of the WRC7 domain, highlighting the coordinating histidine and conserved tryptophan side chains, with labeled protons relevant for NMR analysis. Coordinating histidine 137 (E,H) and Tryptophan 121 (B,C) side-chain resonances were assigned by ^1^H-^1^H TOCSY , ^1^H-^1^H COSY and ^1^H-^15^N long-range HSQC spectra. Observed NOEs confirm interactions consistent with the AlphaFold3 model, validating the predicted side-chain arrangement and structural organization of the WRC domain core (F). ^1^H-^1^H TOCSY and NOESY spectra were acquired in deuterated solvent conditions. Residues are numbered according to GRF7 primary sequence.

**Figure S11.**
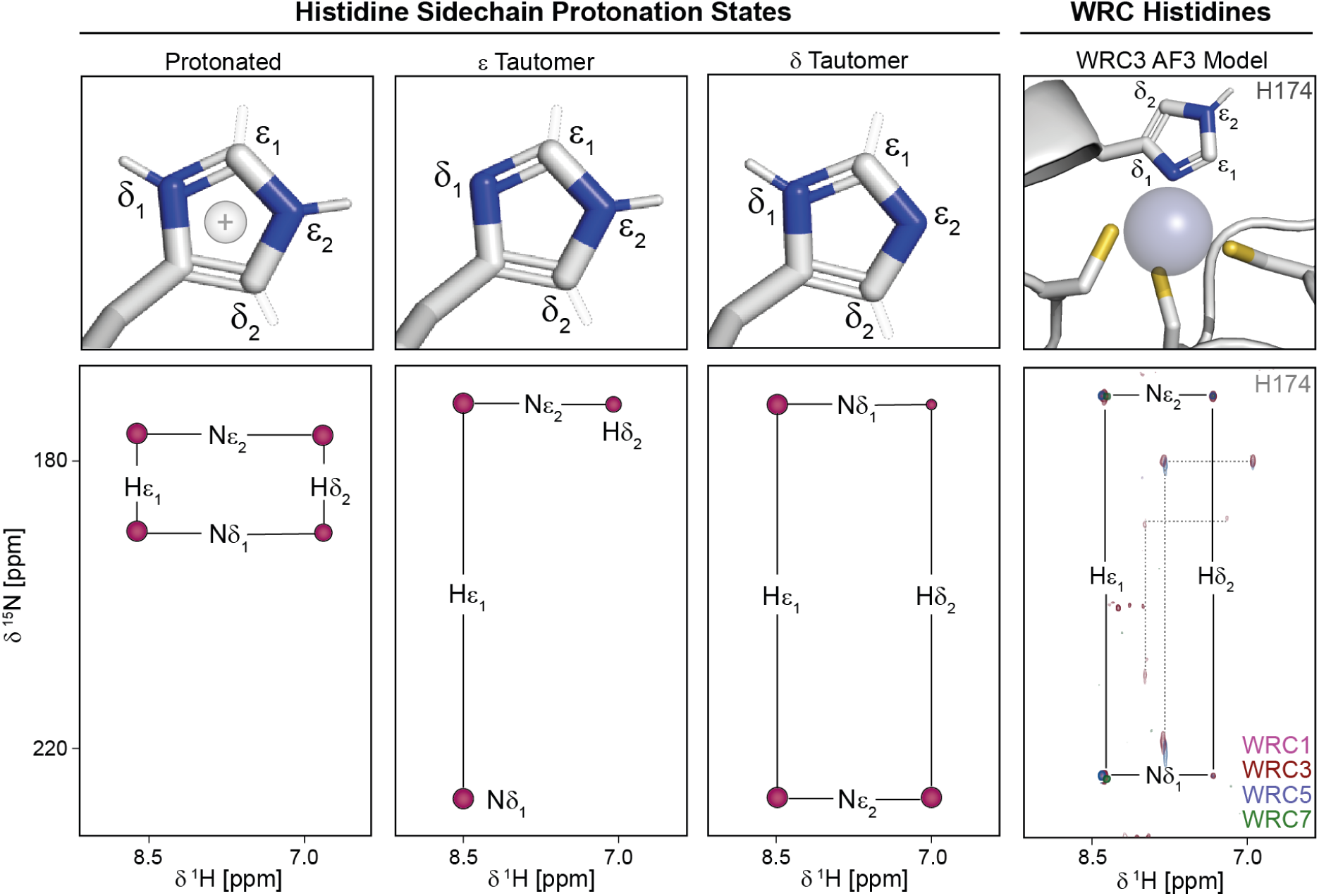
Long-range 1H-15N HSQC spectra of the histidine side-chains of each construct. The left panel shows the expected correlation patterns for histidine residues in three possible tautomeric states (protonated, epsilon and delta) of the imidazole ring. The right panel presents the experimental NMR spectra for all four WRC isoforms (form AtGRF1, AtGRF3, AtGRF5, and AtGRF7). The observed patterns for the coordinating histidines correspond to the epsilon tautomeric state (black lines). Non-coordinating histidine residues are indicated by dashed lines, while coordinating histidines are labeled according to the primary sequence of AtGRF3.

**Figure S12.**
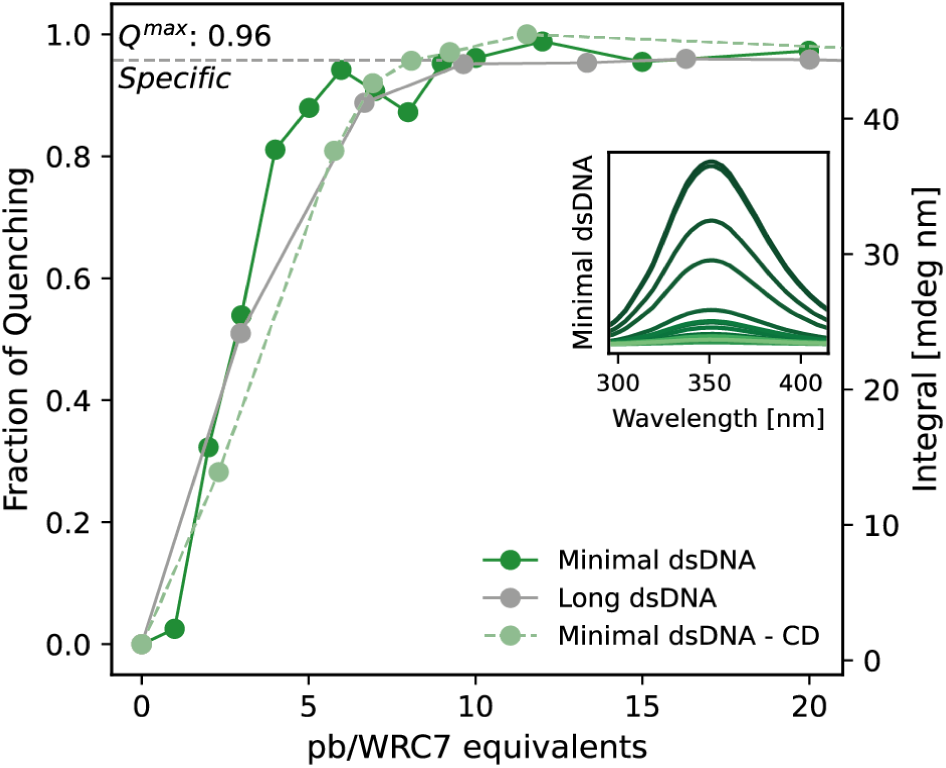
Tryptophan fluorescence quenching of specific minimal (dark green) and long (gray) DNA sequences at different pb/WRC7 equivalents. A similar trend is observed for all probes, indicating the equivalent specificity for the minimal and long specific sequences. Maximal quenching for the long specific probe is indicated by dashed lines. Inset shows the fluorescence emission spectra of the minimal probe for each pb/WRC7 ratio, with darker colors representing lower ratios. The dashed curve (light green) highlights the binding profile derived from the integral CD signal, further validating the consistency and reliability of the tryptophan quenching and CD assays.

**Figure S13.**
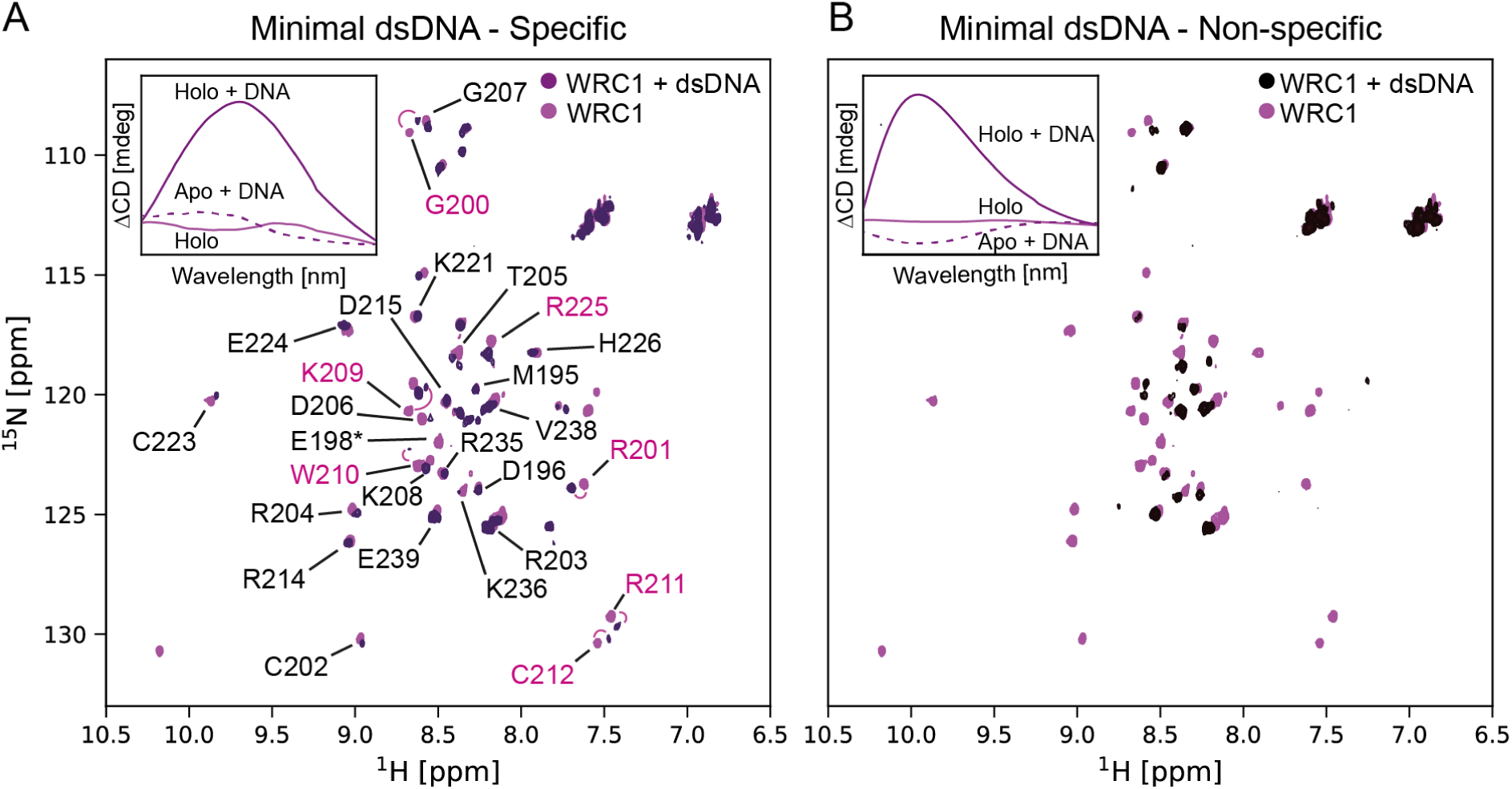
Mapping of DNA-binding residues in the WRC1 domain by NMR spectroscopy and the role of metal coordination in DNA interaction. (A) ^1^H-^15^N HSQC spectrum of WRC1 in the absence (pink) or presence (violet) of specific DNA. Chemical shift perturbation (CSP) analysis reveals residues G200, R201, K209, W210, R211, C212, and R225 as the core DNA recognition interface (pink). The signal corresponding to residue E198 disappears upon DNA addition, suggesting its involvement in DNA binding (asterisk). (B) ^1^H-^15^N HSQC spectrum of WRC1 in the absence (pink) or presence (black) of non-specific DNA. The spectrum shows fewer signals and reduced resolution, indicating the formation of an undefined complex with non-specific DNA. Insets: CD difference spectra of WRC1 in the absence of DNA (holo), in the presence of 0.5 equivalents of DNA (holo + DNA), and in the presence of EDTA and 0.5 equivalents of DNA (apo + DNA) for both minimal probes. The absence of CD signal in the apo + DNA condition indicates the lack of metal coordination, and thus the disruption of the WRC1 structure, prevents DNA recognition and binding.

